# Base editing as a genetic treatment for spinal muscular atrophy

**DOI:** 10.1101/2023.01.20.524978

**Authors:** Christiano R. R. Alves, Leillani L. Ha, Rebecca Yaworski, Cicera R. Lazzarotto, Kathleen A. Christie, Aoife Reilly, Ariane Beauvais, Roman M. Doll, Demitri de la Cruz, Casey A. Maguire, Kathryn J. Swoboda, Shengdar Q. Tsai, Rashmi Kothary, Benjamin P. Kleinstiver

## Abstract

Spinal muscular atrophy (SMA) is a devastating neuromuscular disease caused by mutations in the *SMN1* gene. Despite the development of various therapies, outcomes can remain suboptimal in SMA infants and the duration of such therapies are uncertain. *SMN2* is a paralogous gene that mainly differs from *SMN1* by a C•G-to-T•A transition in exon 7, resulting in the skipping of exon 7 in most *SMN2* transcripts and production of only low levels of survival motor neuron (SMN) protein. Genome editing technologies targeted to the *SMN2* exon 7 mutation could offer a therapeutic strategy to restore SMN protein expression to normal levels irrespective of the patient *SMN1* mutation. Here, we optimized a base editing approach to precisely edit *SMN2*, reverting the exon 7 mutation via an A•T-to-G•C base edit. We tested a range of different adenosine base editors (ABEs) and Cas9 enzymes, resulting in up to 99% intended editing in SMA patient-derived fibroblasts with concomitant increases in *SMN2* exon 7 transcript expression and SMN protein levels. We generated and characterized ABEs fused to high-fidelity Cas9 variants which reduced potential off-target editing. Delivery of these optimized ABEs via dual adeno-associated virus (AAV) vectors resulted in precise *SMN2* editing *in vivo* in an SMA mouse model. This base editing approach to correct *SMN2* should provide a long-lasting genetic treatment for SMA with advantages compared to current nucleic acid, small molecule, or exogenous gene replacement therapies. More broadly, our work highlights the potential of PAMless SpRY base editors to install edits efficiently and safely.

## Introduction

Spinal muscular atrophy (SMA) is a devastating neuromuscular disease that remains a leading cause of infantile death worldwide. SMA is primarily characterized by the death of motor neurons, muscle denervation, and muscle weakness^1,2^. Most SMA cases are caused by loss-of-function mutations within the *Survival Motor Neuron 1* (*SMN1*) gene^1,2^. The most common mutation in *SMN1* is a deletion of exon 7, which leads to abrogation of SMN protein function^3^. An important modifier of SMA severity is the number of copies of a paralogous gene *SMN2*. The sequence of *SMN2* mainly differs from *SMN1* by a synonymous C•G-to-T•A transition in exon 7 (**Fig. 1a**). This C-to-T polymorphism in the 6^th^ nucleotide of *SMN2* exon 7 (henceforth “C6T”; **Fig. 1b**), causes the skipping of exon 7 in most *SMN2* transcripts due to alternative splicing. While *SMN2* still produces ∼10% functional SMN protein, this is not enough to rescue the vast majority of SMA patients^4-6^ (**Fig. 1a**). Targeting *SMN2* transcripts with a small molecule or an antisense oligonucleotide (ASO) to transiently increase the retention of exon 7 demonstrated notable clinical results in infants treated early in the disease process^7-10^.

**Figure 1.**
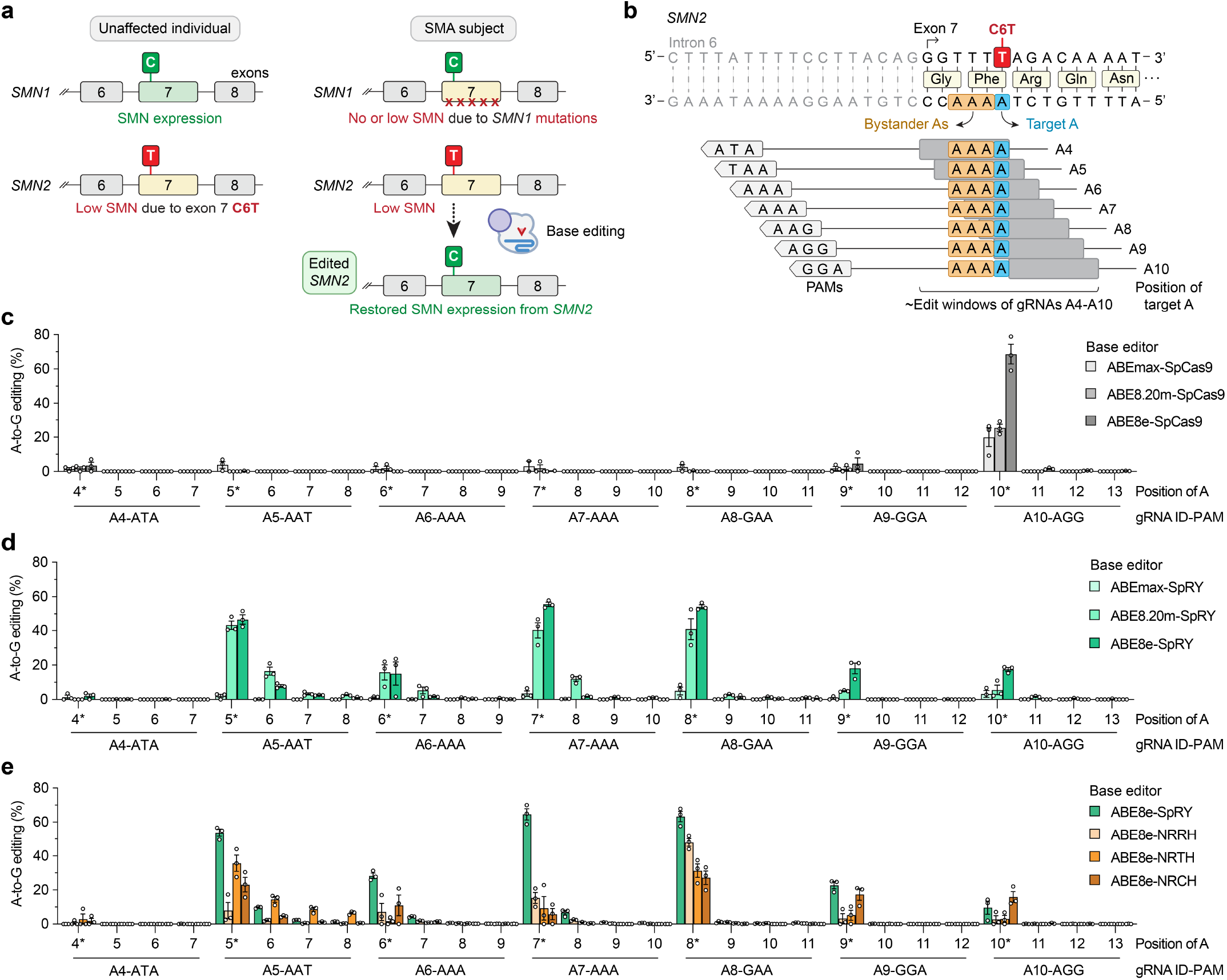
Development of adenine base editing to correct *SMN2* exon 7 C6T. **a**, Schematic of *SMN1* and *SMN2* in unaffected individuals and spinal muscular atrophy (SMA) patients. Mutations in *SMN1* cause SMA due to a depletion of SMN protein, which may be recovered by editing *SMN2*. **b**, Schematic of the *SMN2* exon 7 C-to-T (C6T) polymorphism compared to *SMN1*, with base editor gRNA target sites and their estimated edit windows. **c-d**, A-to-G editing of *SMN2* C6T target adenine and other bystander bases when using ABEs comprised of adenine deaminase domains ABEmax^33,38^, ABE8.20m^35^, and ABE8e^36^ fused to wild-type SpCas9 (**panel c**) or SpRY^37^ (**panel d**), assessed by targeted sequencing. **e**, A-to-G editing of adenines in *SMN2* exon 7 when using SpRY or other relaxed SpCas9 PAM variants^43^, assessed by targeted sequencing. Data in **panels c-e** from experiments in HEK 293T cells; mean, s.e.m., and individual datapoints shown for n = 3 independent biological replicates.

Notwithstanding the development of therapies for SMA, current treatments have limitations and are not a permanent cure. For instance, Risdiplam (Evrysdi, Roche Genentech) is an oral daily administered small molecule splicing modifier that can enhance SMN2 expression by targeting displacement of hnRNP G^11-13^, but this is not a definitive cure for SMA. Nusinersen (Spinraza, Biogen) is an ASO that increases exon 7 inclusion and *SMN2* expression by disabling regulatory elements in *SMN2* intron 7 and is delivered via an intermittent regimen of intrathecal injections across the lifespan of the patient^8,9,14^. Because Nusinersen is injected into the spinal fluid, it is not expected to modify *SMN2* expression in peripheral tissues that may play a role in SMA^15-18^. Likewise, exogenous gene addition using an adeno-associated virus (AAV) vector expressing *SMN1* such as onasemnogene abeparvovec (Zolgensma, Novartis) presents many challenges^19^, including unknown longevity of expression from the AAV transgene^20^, and eventual decay of efficacy in dividing cells due to AAV dilution^21^. Moreover, persistent supraphysiological expression of *SMN1* from ubiquitous promoters may cause toxicity^22^. Although the success of approved therapies has minimized the previously high infantile and childhood morbidity and mortality associated with SMA, their emerging limitations underscores the need to further improve upon existing therapies in terms of the scope and durability of SMN protein expression. Thus, there is an unmet need to develop single-dose therapies that permanently increase SMN levels.

Genome editing technologies capable of permanently editing *SMN2* to restore SMN levels could overcome several of these challenges (**Fig. 1a**). The pursuit of genome editing methods to treat the diverse spectrum of *SMN1* mutations is less feasible since it would necessitate patient- and mutation-specific optimization of a variety of editing approaches. Instead, the development of a strategy to directly revert the *SMN2* C6T polymorphism could prove to be a broadly applicable editing strategy to increase SMN expression for SMA patients. This contrasts with other editing approaches that have been explored to treat SMA, including CRISPR-Cas9 or Cas12a nucleases to attempt homology-directed repair of *SMN2*, or the use of nucleases to modify intronic splicing regulatory elements (SREs) in intron 7 ^23-25^ (which are also the target of nusinersen^26^). However, due to the number of copies and high similarity between the *SMN1* and *SMN2* genes, nuclease-based approaches that create DNA double-strand breaks (DSBs) carry the risk of creating large chromosomal deletions, translocations, or other undesirable DSB-related consequences^27-31^.

The optimization of DSB-independent strategies would therefore offer advantages over nuclease-based strategies by reducing the likelihood of unwanted genome-scale changes and potentially improving editing efficiency. Base editors (BEs) are one potential technology capable of installing point mutations without intentionally creating DNA DSBs^32^. Typically, BEs are composed of a fusion of a CRISPR-Cas enzyme (i.e. *Streptococcus pyogenes* Cas9; SpCas9) to a deaminase domain, and when directed by a guide RNA (gRNA) to genomic sites, BE complexes can initiate edits of specific DNA bases. Adenine base editors (ABEs) catalyze A•T to G•C edits^32-34^ using an evolved TadA deaminase to convert adenines to inosines, which are read as guanines by polymerases^33^. However, the deaminase domain can only act in a narrow ‘edit window’ within the gRNA-Cas target site at a fixed distance from the protospacer-adjacent motif (PAM; **Fig. 1b**). Thus, the efficiency of base editing is dependent on the availability of Cas9 variant enzymes that can recognize a range of PAMs to maximize editing of the intended base while minimizing editing of unwanted nearby bases (so-called bystander edits). We therefore hypothesized that the identification and optimization of ABEs capable of editing only the target C6T adenine in *SMN2* exon 7 could correct the transition mutation and restore SMN expression (**Fig. 1a**).

Here we explored the potential of various ABE-based strategies to treat SMA, including correction of C6T in *SMN2* and modification of *SMN2* intron 7 SREs. We identified efficient and precise combinations of ABEs and gRNAs capable of specific A-to-G editing in HEK 293T cells with low levels of bystander editing. We then extended our *SMN2* C6T editing approach into SMA patient-derived fibroblasts, leading to increased exon 7 retention in *SMN2* transcripts and elevated expression of SMN protein. We assessed the genome-wide safety of this approach and while we did not observe unwanted off-target editing in fibroblasts, off-target editing was reduced in HEK 293T cells when using high-fidelity variants. Finally, we demonstrate the feasibility of translating this approach *in vivo* in a mouse model of SMA via AAV-mediated delivery of the ABE and gRNA. Taken together, our results demonstrate that ABE-mediated editing of *SMN2* C6T leads to substantial increases in SMN protein levels, establishing the potential of a new therapeutic approach to treat SMA.

## Results

### Development of ABEs to edit *SMN2* C6T

We first explored whether ABEs could correct the C•G-to-T•A C6T transition in *SMN2* exon 7 by performing experiments in HEK 293T cells. There are two potential challenges for this approach. First, there is only one gRNA target site with an NGG PAM accessible with wild-type (WT) SpCas9, which positions the target adenine in position 10 of the spacer (A10; **Fig. 1b**). The 10^th^ nucleotide of the spacer is near the border of the canonical ABE edit window, although engineered adenine deaminase domains have expanded editing potency across a wider sequence space^35,36^. A second potential complicating factor is that the target adenine is bordered by three additional adenines (**Fig. 1b**), which may lead to unwanted bystander edits of unknown functional consequences. Ideally the ABE would edit only the target adenine with minimal editing of the neighboring adenines. To identify an efficient and precise ABE, we designed seven different gRNAs against sites harboring various PAMs, tiling the edit window of the ABEs by placing the target base at positions A4-A10 (**Fig. 1b**). Due to the availability of only one site with an NGG PAM, in addition to performing these experiments with WT SpCas9 (**Fig. 1c**), we also utilized our previously engineered SpCas9 variant that has a relaxed tolerance for PAMs^37^, SpRY (**Fig. 1d**).

In our initial experiments using ABEmax^33,38^, as expected, WT SpCas9 showed measurable activity only when using the A10 gRNA that targets the site encoding an NGG PAM (**Fig. 1c**). Conversely, ABEmax-SpRY exhibited A-to-G editing using nearly every gRNA, although with modest editing (<5%; **Fig. 1d**). We therefore explored the use of two engineered ABE domains previously shown to improve on-target editing (ABE8.20m and ABE8e)^35,36^. Using these two additional ABE domains, WT SpCas9 and SpRY ABEs mediated substantially higher levels of A-to-G editing (**Figs. 1c** and **1d**, respectively). With WT SpCas9 ABEs, once again only gRNA A10 was conducive to high levels of editing (**Fig. 1c**). When using SpRY ABEs, we observed high levels of editing using gRNAs A5 (NAT PAM), A7 (NAA PAM), and A8 (NAA PAM) (**Fig. 1d**). In general, ABE8e-based enzymes led to higher levels of editing compared to ABE8.20m (**Figs. 1c** and **1d**). Analysis of different 5’ gRNA spacer architectures ^39-41^ did not substantially impact editing efficiency (**Sup. Note 1** and **Sup Figs. 1a** and **1b**).

We found that while high levels of A-to-G editing could be achieved at the intended target adenine (*SMN2* C6T), bystander editing of the three neighboring adenines was generally minimal (**Figs. 1c** and **1d**). When using ABE8e-WT with gRNA A10 or ABE8e-SpRY with gRNA A8, we observed the highest levels of A-to-G editing with near-background levels of bystander editing (**Fig. 1d**). The lack of bystander editing of the adjacent adenines is partially supported by previous reports^42^ of preceding adenines being inhibitory to ABE efficiency (as is the case of each of the 3 bystander As in these target sites), while preceding thymines promote ABE editing (as with the target adenine). However, additional transfections using various ABEs and up to 7 other gRNAs targeting sites with poly-adenine stretches did not reveal as striking enrichment for editing at the 5’ adenine in all cases (**Sup. Figs. 2a-2f** and **Sup. Note 2**).

**Figure 2.**
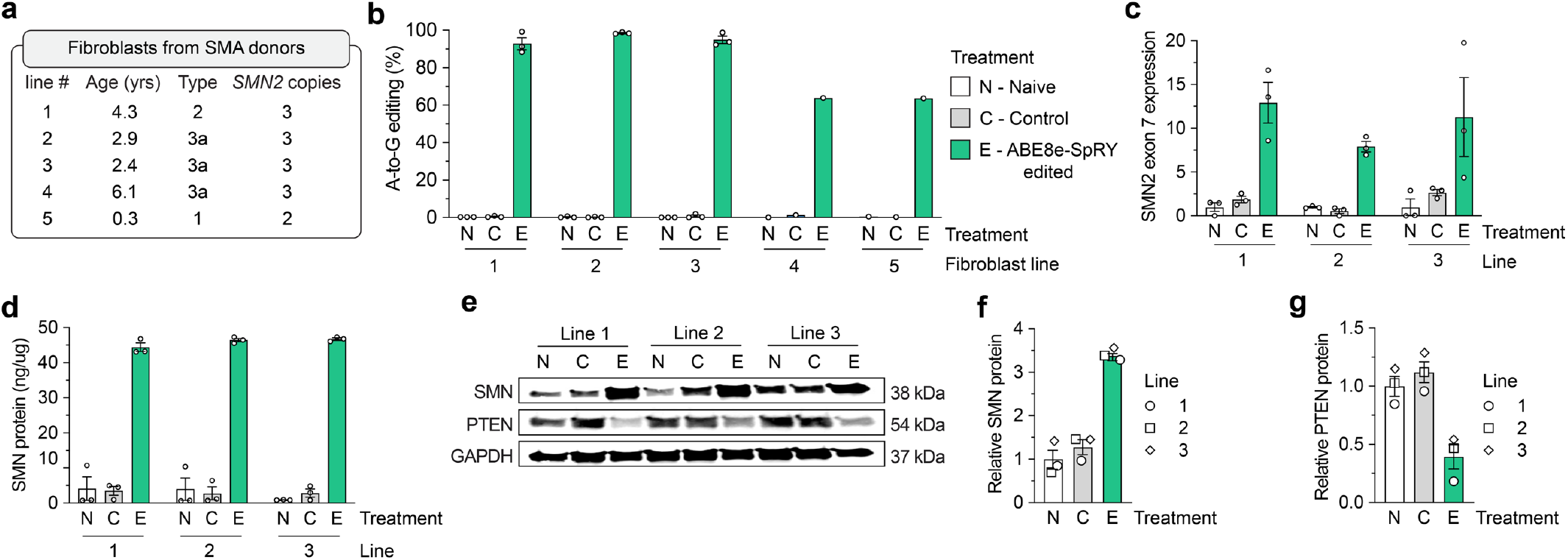
*SMN2* C6T editing and phenotypes in SMA patient-derived fibroblasts. **a**, Characteristics of five different SMA donors; all lines harbor a homozygous deletion of exon 7 in *SMN1*. **b**, A-to-G editing of the C6T adenine in *SMN2* exon 7 across five SMA fibroblast cell lines transfected with ABE8e-SpRY and gRNA A8, assessed by targeted sequencing. Naïve (N) cells were untransfected; Control (C) cells were treated with ABE8e-SpRY and a non-targeting gRNA. **c**, SMN2 exon 7 mRNA expression across three edited (E) SMA fibroblast lines, measured by ddPCR. Transcript levels normalized by GAPDH mRNA. **d**, SMN protein levels determined by an SMN-specific enzyme-linked immunosorbent assay (ELISA). **e**, Representative immunoblot for SMN, PTEN, and GAPDH protein levels across Naïve, Control, or ABE8e-SpRY treated SMA fibroblast lines. **f**,**g**, Quantification of SMN and PTEN (**panels f** and **g**, respectively) protein levels normalized to GAPDH and the Naïve treatment, determined by immunoblotting. For all assays, GFP-positive fibroblasts were sorted post-transfection and grown in for at least 3 passages; samples from three independent passages were collected for lines 1, 2 and 3 (passages 4-6; see **Sup. Fig. 6a**), and one passage was collected for lines 4 and 5. For **panels b-d, f**, and **g**, mean, s.e.m., and individual datapoints shown for n = 3 independent biological replicates from separate passages (unless otherwise indicated).

We also assessed editing with other previously described SpCas9 PAM variants, including SpG (capable of targeting sites with NGN PAMs)^37^ and other relaxed PAM variants SpCas9-NRRH, NRTH, and NRCH (which can target sites with their namesake PAMs; R is A or G, and H is A, C, or T)^43^. For these experiments we utilized ABE8e due to its superior A-to-G editing efficiency. With ABE8e-SpG paired with gRNA A9 (NGA PAM) or gRNA A10 (NGG), we observed modest levels of on-target editing (**Sup. Fig. 3**) at efficiencies lower than ABE8e-WT and gRNA A10 (**Fig. 1c**) or ABE8e-SpRY with gRNAs A5, A7, or A8 (**Fig. 1d**). With ABE8e-NRRH, -NRTH, and -NRCH, we observed consistently lower on-target editing compared to ABE8e-SpRY (**Fig. 1e**).

**Figure 3.**
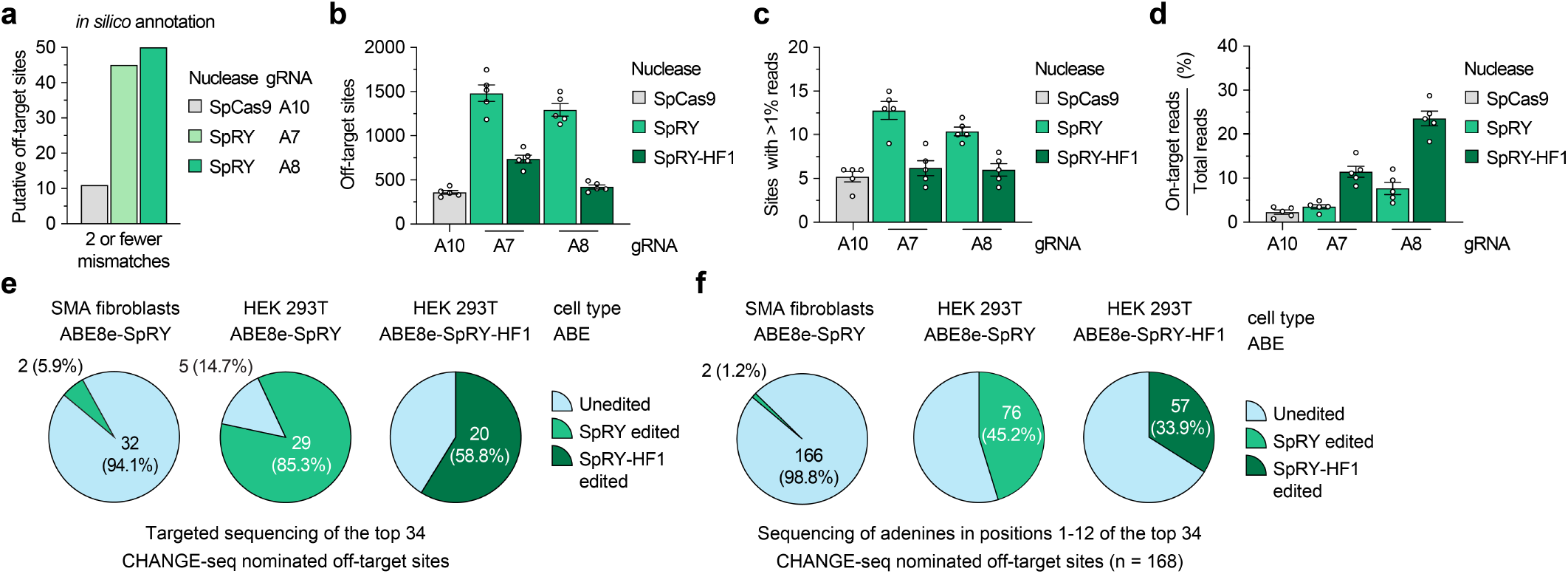
Analysis of *SMN2* C6T base editing specificity. **a**, Number of putative off-target sites in the human genome with up to 2 mismatches for each gRNA, annotated by CasOFFinder^53^. Predictions for SpCas9 utilized an NGG, NAG, or NGA PAM; with SpRY, a PAMless NNN search. **b**, Total number of CHANGE-seq detected off-target sites, irrespective of assay sequencing depth. **c**, Number of CHANGE-seq identified off-target sites that account for greater than 1% of total CHANGE-seq reads and are common across all 5 SMA fibroblast lines. **d**, Percentage of CHANGE-seq reads detected at the on-target site relative to the total number of reads in each experiment. For **panels b, c**, and **d**, mean, s.e.m, and individual datapoints shown for n = 5 independent biological replicate CHANGE-seq experiments (performed using genomic DNA from each of the 5 SMA fibroblast lines). **e**,**f**, Summary of targeted sequencing results from ABE-edited SMA fibroblasts or HEK 293T cells at the top 34 CHANGE-seq nominated off-target sites (common sites across all 5 SMA fibroblast lines and treatments with SpRY or SpRY-HF1), analyzing statistically significant editing of any adenine in the target site (**panel e**) or of all adenines in positions 1-12 of each of the 34 target sites (**panel f**).

Given that the number of genomic sites encountered by Cas9 enzymes with minimal PAM requirements is expanded, one consideration when using relaxed PAM variants is the potential to observe editing at an increased number off-target sites^37,43^. To mitigate potential genome-wide off-target editing, we determined whether the ABE8e constructs were compatible with two previously described high-fidelity SpCas9 variants that eliminate or minimize off-target editing^44,45^. We generated ABE8e fusions to SpCas9 and SpRY in the presence of HF1 substitutions that were previously shown to eliminate nearly all off-target edits^45^, and the HiFi mutation that was previously shown to reduce levels of off-target editing^44^. The HF1 and HiFi versions of ABE8e-SpCas9 exhibited a substantial loss in on-target *SMN2* editing (**Sup. Fig. 4a**). Conversely, in some cases the HF1 and HiFi variants of ABE8e-SpRY retained high levels of editing (albeit at levels lower than ABE8e-SpRY; **Sup. Fig. 4b**), potentially indicating that the high-fidelity mutations may exhibit a greater loss in on-target base editing when the target adenine is closer to the boundary of the ABE edit window. We observed similar levels of bystander editing for all conditions (**Sup. Figs. 4a** and **4b**), and a minor impact of the 5’ gRNA spacer architecture on the efficiency of high-fidelity variants (**Sup. Note 1** and **Sup. Figs. 4c-4f**).

**Figure 4.**
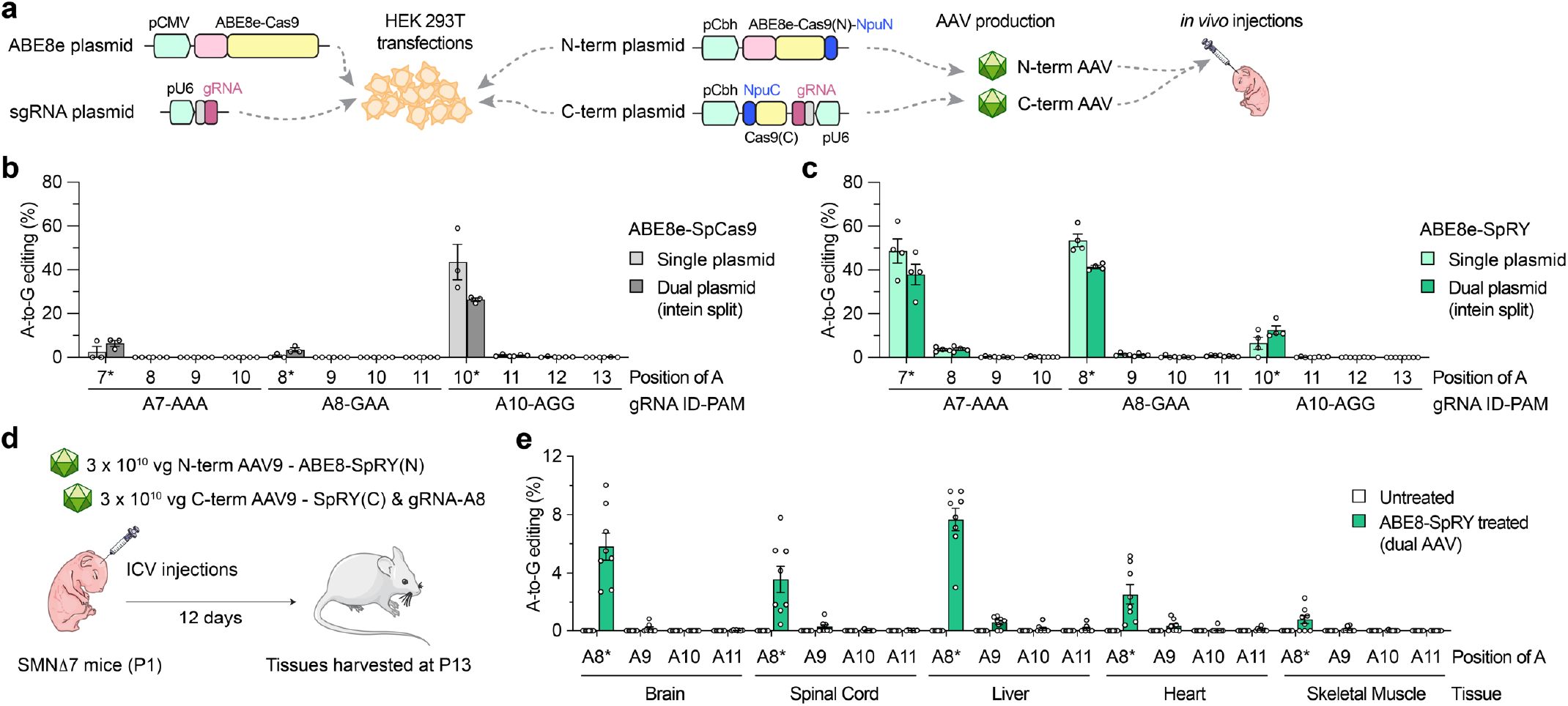
AAV-mediated delivery of base editors for *in vivo SMN2* C6T editing. **a**, Schematics of conventional and intein-split plasmids for ABE and gRNA delivery in cells and *in vivo*. gRNA, guide RNA; NpuN/NpuC, N- and C-terminal intein domains; Cas9(N) and Cas9(C), N- and C-terminal fragments of SpCas9 variants. **b**,**c**, A-to-G editing of *SMN2* C6T target adenine and other bystander adenines when using ABE8e-SpCas9 with gRNA A10 (**panel b**) or ABE8e-SpRY with gRNA A8 (**panel c**), assessed by targeted sequencing. Data in **panels b** and **c** from experiments in HEK 293T cells; mean, s.e.m., and individual datapoints shown for n = 3 independent biological replicates. **d**, Schematic of P1 intracerebroventricular (ICV) injections in SMNΔ7 mice with dual AAV9 vectors that express intein-split ABE8e-SpRY and gRNA-A8. **e**, A-to-G editing of *SMN2* exon 7 adenines following ICV injections of AAV encoding ABE8e-SpRY with gRNA A8 (**panel d**). Editing across various tissues (without sorting for transduced cells) assessed by targeted sequencing of n = 6 treated and n = 8 untreated (sham injection) SMNΔ7 mice; mean, s.e.m., and individual datapoints shown.

Taken together, these results demonstrate that optimized ABEs are selective for editing the target C6T adenine in *SMN2* exon 7 with low-level bystander editing, particularly when utilizing ABE8e-WT with gRNA A10 or ABE8e-SpRY with gRNA A8. Given that both enzyme-gRNA pairs exhibited comparable on-target editing of the target adenine, we selected the ABE8e-SpRY and gRNA A8 pair for further study due to a more optimal positioning of the target adenine in the ABE edit window and its enhanced compatibility with high-fidelity variants.

### Simultaneous base editing of *SMN2* exon 7 and intronic splicing silencers in human cells

*SMN2* exon 7 alternative splicing is caused by the C6T mutation that abrogates *SMN2* exon 7 inclusion through multiple mechanisms, including disruption of binding sites for pre-mRNA-splicing factors SF2/ASF^46-48^, and creation of binding sites for the nuclear ribonucleoproteins hnRNPA1 and hnRNPA2 that repress exon 7 inclusion in *SMN2* mRNA^48-50^ (**Sup. Fig. 5a**). Interestingly, there are two intronic splicing silencer (ISS) binding sites for these ribonucleoproteins in intron 7 of *SMN2* (ISS-N1 and ISS+100; **Sup. Fig. 5a**). Disruption of these ISSs via genome editing could be an alternate approach to increase *SMN2* exon 7 inclusion and SMN expression. Indeed, nuclease-mediated knockout of these two ISSs using CRISPR-Cas12a enzymes was shown to enhance SMN protein expression from *SMN2*^24^. We therefore explored the use of ABE8e-SpRY to generate A-to-G edits within the ISSs, which would be compatible with our C6T editing approach and would hold advantages over nuclease-mediated editing given substantially reduced levels of DNA DSBs.

To determine whether ABE8e-SpRY could edit the ISS-N1 and ISS+100 regulatory elements in *SMN2*, we designed 12 gRNAs that would position the edit window of ABE8e-SpRY over adenine bases of ISS-N1 and ISS+100 (**Sup. Fig. 5b**). We transfected HEK 293T cells using ABE8e-SpRY and each of the 12 ISS-targeting gRNAs in the absence or presence of the gRNA A8 (for simultaneous targeting of exon 7 C6T). When targeting either ISS, we observed editing of various adenines, albeit at efficiencies lower than C6T targeting (**Sup. Figs. 5c** and **5d**; **Sup. Note 3**). We then explored multiplex targeting of *SMN2* by co-transfecting gRNA A8 for C6T and the ISS gRNAs, which led to efficient C6T editing (**Sup. Figs. 5e** and **5f**) and maintenance of ISS-N1 and ISS+100 editing (**Sup. Figs. 5c** and **5d**; **Sup. Note 3**). These results demonstrate that a dual-editing approach minimally impacts *SMN2* C6T editing and could be explored in future studies as a multiplex strategy to increase the likelihood of exon 7 inclusion in the *SMN2* transcript, with minimal levels of DSBs.

### Assessment of phenotypic changes in ABE-edited SMA patient-derived fibroblasts

We then assessed the efficiency of our ABE approach in five SMA patient-derived fibroblast cell lines (**Fig. 2a**). We transfected plasmids encoding ABE8e-SpRY-P2A-EGFP and gRNA A8 to permit sorting for GFP+ fibroblasts (**Sup. Fig. 6a**). Analysis of editing in these sorted cells revealed high levels of *SMN2* C6T editing across all five SMA cell lines compared to naïve samples that were untransfected or control samples transfected with SpRY-ABE8e and a non-targeting gRNA (**Fig. 2b**). For three fibroblast lines, we observed near complete editing (>90% A-to-G edits), while for two additional lines we observed A-to-G editing around 60%. Importantly, there was little (<1%) bystander editing of adjacent adenines across all five fibroblast lines (**Sup. Fig. 6b**). These results demonstrate that our base editing strategy for C6T is extensible to, and efficient in, SMA patient-derived cells.

Next, we sought to assess the extent to which ABE-mediated editing of *SMN2* C6T would increase SMN transcript and protein levels using the three nearly completely edited SMA fibroblast lines (1, 2, and 3). At the transcript level, we observed a ∼6.3-fold mean increase in *SMN2* exon 7 mRNA expression compared to control cells (**Fig. 2c**; from *SMN2* transcripts alone since each fibroblast line harbors homozygous deletions for *SMN1* exon 7). We also detected a ∼2.7-fold increase in expression of the early *SMN* transcript (across the junction of exons 1 and 2 for *SMN1* and *SMN2*) (**Sup. Figs. 6c** and **6d**), suggesting increased stability of full-length *SMN* mRNA in ABE-edited SMA-fibroblasts.

We investigated concomitant alterations in SMN protein levels in the edited fibroblasts and observed a substantial ∼15-fold mean increase in SMN protein levels when compared with control cells as measured by an ELISA (**Fig. 2d**). Moreover, immunoblotting revealed increased SMN protein in edited fibroblasts (**Figs. 2e-f**), although with a lower magnitude than observed with the ELISA (**Fig. 2d**). Interestingly, restoring SMN protein was associated with a decrease in PTEN protein levels (**Figs. 2e** and **2g**). Elevated PTEN in SMA cells has previously been associated with increased apoptosis^51,52^, indicating that *SMN2* C6T editing may lead to reduced PTEN and improved viability of ABE-edited SMA cells. Together, these functional assays demonstrate that precise editing of *SMN2* C6T in SMA patient-derived fibroblasts results in consistently increased full-length SMN transcript and protein levels.

### Off-target nomination and validation

To identify and evaluate potential off-target sites, we utilized computational and experimental methods to profile our Cas9 and gRNA combinations that exhibited the highest on-target *SMN2* C6T editing. Comprehensive *in silico* off-target annotation using CasOFFinder^53^ for WT SpCas9 with gRNA A10 (using NGG, NAG, or NGA PAMs^54-56^), and SpRY with gRNAs A5, A7, and A8 (using a PAMless NNN search) resulted in 11, 94, 45, and 50 off-targets with 2 or fewer mismatches, including the off-by-one mismatch off-target of *SMN1* (**Fig. 3a** and **Sup. Figs. 7a**-**7c**).

We then performed an unbiased biochemical off-target nomination assay, CHANGE-seq^57^, using the gDNA from each SMA-fibroblast line used in this study to experimentally nominate putative genome-wide off-targets. CHANGE-seq experiments were performed using WT SpCas9 nuclease with gRNA A10 (**Sup. Figs. 8** and **9**) and SpRY and SpRY-HF1 nucleases^37,45^ with gRNA A7 (**Sup. Figs. 10** and **11**) and gRNA A8 (**Sup. Figs. 12** and **13**). The SpRY-HF1 variant includes fidelity-enhancing mutations previously shown to eliminate nearly all off-target edits with nucleases in cells^37,45^. CHANGE-seq identified a range of off-targets for each nuclease and gRNA combination, with SpRY leading to the highest number compared to WT SpCas9 or SpRY-HF1 (**Fig. 3b, Sup. Figs. 8**-**13**, and **Sup. Note 4**). We observed a comparable number of off-target sites common across each of the five fibroblast lines for SpRY, SpRY-HF1, and SpCas9 treatments, including those that reached greater than 1% of total CHANGE-seq read counts in each experiment (**Fig. 3c** and **Sup. Note 4**). The off-by-one mismatch *SMN1* off-target was not detected due to homozygous loss in all five fibroblast lines. The intended *SMN2* on-target site was typically the most efficient and abundant CHANGE-seq site detected for SpRY and SpRY-HF1 treated samples (**Fig. 3d** and **Sup. Figs. 10**-**13**); in most cases, the on-target site for WT SpCas9 with gRNA A10 was targeted less efficiently than several off-targets, resulting in a smaller fraction of total CHANGE-seq reads assigned to the on-target site (**Fig. 3d** and **Sup. Figs. 8**-**9**). However, as expected due to its expanded PAM tolerance, SpRY with gRNA A7 or A8 resulted in a larger number of total CHANGE-seq detected off-target sites compared to WT SpCas9, although most were detected at much lower efficiencies compared to the on-target sites (**Fig. 3b, Sup. Figs. 8**-**13, Sup. Note 4**, and **Supplementary Table 1**). Use of SpRY-HF1 substantially reduced the number of CHANGE-seq detected off-targets and led to an enriched targeting of the *SMN2* C6T on-target site, as compared to SpRY treatments (**Figs. 3b-3d**; **Sup. Figs. 10**-**13**).

To assess potential off-target editing in cells, we performed targeted sequencing using genomic DNA from SMA-fibroblasts treated with ABE8e-SpRY and gRNA A8. We assessed the top 34 CHANGE-seq detected off-target sites that were consistently identified across the five SMA-fibroblast lines treated with either SpRY or SpRY-HF1 using gRNA A8 (**Supplementary Table 1**). Targeted sequencing across 3 different fibroblast lines revealed high levels of editing at the C6T on-target site compared to naïve cells (**Sup. Fig. 14a**), consistent with our prior results. Analysis of off-target editing at adenines in positions 1-12 of the spacers of the 34 off-targets in edited SMA fibroblasts revealed background-levels of editing at all but two adenines in two off-target sites (**Figs. 3e** and **3f**, and **Supplementary Table 2**). These results demonstrate that high specificity editing can be achieved when using near PAMless SpRY-based ABEs in fibroblasts, similar to a previous report that utilized an engineered SpCas9 variant with an altered but more specific PAM preference^58^.

We also sequenced the on- and off-target sites of genomic DNA from HEK 293T cells treated with ABE8e-SpRY or ABE8e-SpRY-HF1 and gRNA A8 (**Sup. Fig. 14b**). Analysis revealed detectable off-target editing in at least one adenine in 29 of 34 off-target sites when using ABE8e-SpRY or 20/34 with ABE8e-SpRY-HF1 (**Fig. 3e**), potentially reflecting a difference in the transfection efficiencies and expression levels of the enzymes between sorted fibroblasts and HEK 293T cells. Again, in many cases, the HF1 variant reduced the number of or efficiency of editing at off-target adenines (**Figs. 3e** and **3f**, and **Sup. Fig. 14b**). In summary, these findings demonstrate that the off-target profile of our ABE-mediated C6T edit is highly specific in fibroblasts but varies depending on the cell-type and/or transfection approach.

### *In vivo* base editing in a mouse model of SMA

Given our cell-based results indicating efficient C6T editing using ABE8e-SpRY, we sought to assess the efficacy of *in vivo* editing in a mouse model of SMA. To treat SMA, the primary target tissue for C6T editing is motor neurons in the central nervous system (CNS), although previous reports indicate the importance of SMN expression in peripheral tissues^15-18^. For *in vivo* delivery of ABE8e-SpRY we utilized an intein-mediated dual-AAV approach due to the size of the construct, splitting the editing components between two vectors encoding TadA8e fused to an N-terminal fragment of Cas9, and the C-terminal fragment of Cas9 with the gRNA (similar to as previously described^58,59^) (**Figs. 4a** and **Sup. Figs. 15a**-**15b**). Prior to generating the AAV, we transfected the AAV plasmids encoding the split ABEs into HEK 293T cells and analyzed on-target editing of *SMN2* C6T. The split WT ABE8e-SpCas9 with gRNA A10 resulted in ∼30% on-target A-to-G editing, levels lower than what we observed for the conventional ABE construct (**Fig. 4b**). However, when assessing editing with the split ABE8e-SpRY constructs using either gRNA A7 or A8, we observed approximately ∼40% editing, levels more comparable to the single plasmid ABE (**Fig. 4c**).

We then performed a pilot study in neonatal P1 mice to compare the CNS transduction efficiency of two different AAV capsids, AAV9 and AAV-F^60^ after intracerebroventricular (ICV) injections (**Sup. Figs. 16a**-**16e** and **Sup. Note 5**). Since we observed comparable spinal cord transduction between either serotype and given that AAV9 is the serotype of the FDA approved Zolgensma gene therapy to treat SMA^19^, we selected AAV9 for *in vivo* studies.

To determine whether we could edit *SMN2* C6T using ABE8e-SpRY and gRNA A8 *in* vivo, we selected SMNΔ7 mice as a model for experiments (**Sup. Note 6**). SMNΔ7 mice are transgenic for the human *SMN2* gene and SMNΔ7 cDNA and have a severe and rapid phenotype under *Smn*^-/-^ conditions^61-63^. We performed ICV injections of 3 × 10^10^ vector genomes (vg) of each of the N- and C-terminal AAVs in P1 SMNΔ7 neonatal mice, and we harvested tissues from mice at P13 for sequence analysis (**Fig. 4d** and **Sup. Note 6**). Whether from the Smn^+/+^, Smn^+/-^, or Smn^-/-^ genotypes (**Sup. Fig. 17a**), we observed on-target (*SMN2* C6T) A-to-G editing in multiple tissues, including the brain, spinal cord, liver, heart, and skeletal muscle (**Fig. 4e** and **Sup. Fig. 17b-c**). In the tissues of primary interest, we detected ∼4% (spinal cord) and ∼6% (brain) A-to-G editing on average (range of 2% to 10% across different mice) with low levels of bystander editing in these edited tissues (**Fig. 4e** and **Sup. Note 6**). These levels of editing in bulk tissue are similar to other studies that delivered base editors via ICV injections in neonatal mice via AAV vectors or eVLPs^59,64^, despite differences in experimental schemes. Since the lifespan of this SMA mouse model is typically less than 15 days^65^, there was no obvious improvement in SMA mice phenotype (**Sup. Note 6**) and future studies may be necessary to extend SMA mouse survival to allow the investigation of longer-term base editing effects. Together, these results reveal that the delivery of this optimized ABE via a dual-AAV vector approach results in *SMN2* C6T editing *in vivo* in an SMA mouse model.

## Discussion

SMA is a progressive neuromuscular disease caused by mutations in the *SMN1* gene that remains a leading cause of infantile death worldwide. Type 1 SMA is the most frequent clinical subtype accounting for more than 50% of cases, characterized by severe denervation that causes untreated patients to die early in life^66,67^. Fortunately, extensive investigation into SMA therapeutics has culminated in FDA approved medicines with striking benefits for patients (risdiplam, nusinersen, and onasemnogene abeparvovec). These therapies have profoundly altered the trajectory of newborns with SMA. In spite of this, each therapy has limitations including strict dosing regimens and open questions surrounding toxicity and long-term efficacy, that together underscore and motivate the need for continued advances in genetic therapies. Early indications from clinical trials utilizing genome editing technologies to treat other diseases indicate that one-time persistent treatments may be within reach^68,69^.

Here we describe the optimization of pan-SMA mutation base editing approaches within the paralogous *SMN2* gene irrespective of the patient *SMN1* mutation. By leveraging our previously described broad targeting enzyme SpRY, we identified a set of BEs and gRNAs capable of positioning the BE edit window over the *SMN2* exon 7 C6T base, leading to high editing efficiencies in HEK 293T cells and SMA patient-derived fibroblasts. Specific C6T editing led to restoration of *SMN2* mRNA and protein expression, and this strategy was translatable to *in vivo* experiments in mice via dual AAV vector delivery. Importantly, we observed minimal unwanted bystander or genome-wide edits in fibroblasts (and also demonstrated that improved specificity variants can reduce levels of off-target editing), suggesting that PAMless BEs can be target-specific and that their flexibility offers advantages for treating a range of diseases.

We detailed two distinct base editing approaches to modify *SMN2*, including the installation of a specific edit within exon 7 C6T, or the disruption of two ISSs within intron 7. These approaches offer several advantages over prior genome editing pursuits. The design of *SMN2* editing strategies capable of treating all SMA patients obviates the necessity of optimizing patient-specific medicines for the individual *SMN1* mutations or deletions. Base editing approaches that are not reliant on nucleases should avoid unwanted DSB-related consequences^27-31^. A previously study^24^ described ISSs knockout methods that could potentially introduce large deletions (>600 kb) between the *SMN1* and *SMN2* genes in patients (who are not *SMN1*-null), or between duplicated *SMN2* copies on the same chromosome. Notably, our ISS-targeting strategy with BEs capitalizes on the same mechanism targeted by the FDA-approved ASO nusinersen, suggesting that our proof-of-concept editing experiments should enhance SMN protein levels as previously shown using nucleases^24^. Beyond the two approaches we outline herein, additional splicing-regulatory elements including potential ISSs were recently described that may also serve as appropriate base editing targets^70^.

Via dual AAV-vector delivery of ABE8e-SpRY, we achieved editing in the primary target tissues for reversing SMA pathology including spinal cord and brain. The *in vivo* editing efficiency in bulk tissues (i.e. homogenized P13 brain) was similar to other studies that delivered base editors via ICV injections in neonatal mice via AAV vectors or engineered virus-like particles (eVLPs)^59,64^ with analysis of editing at later timepoints (3-4 weeks vs. P13 for our experiments). Future experiments that assess alternate delivery vehicles (i.e. AAV vectors with improved CNS transduction^71-73^, eVLPs, or nanoparticles), additional injection paradigms or routes (i.e. *in utero*^*74*^), or longer time points may lead to higher levels of editing in both the CNS and peripheral tissues. Furthermore, alternate approaches may also provide an extended therapeutic window in the severe SMNΔ7 mouse model for expression of the ABE8e-SpRY to perform C6T editing prior to irreversible pathology. For instance, the use of other SMA models and the exploration of co-therapies using small molecules or ASOs^15,75-77^ may enable enhanced base editing and phenotypic recovery *in vivo*. Finally, the continued development of improved genome editing technologies, including those capable of more efficient and specific editing at non-canonical PAMs^56,78^, may improve the efficacy and safety of targeting.

In summary, we develop a base editing approach to treat SMA by modifying *SMN2* to increase SMN expression. Compared to other approved SMA therapies, genome editing technologies offer hope for a long-lasting therapeutic effect. More broadly, our work underscores how highly versatile CRISPR enzymes like SpRY can be leveraged to create specific genetic edits in model organisms^79^, establishing a blueprint to extend this approach to additional neuromuscular disorders and other classes of disease.

## Acknowledgements

We acknowledge the following individuals: C. Tou (ddPCR), E. King (AAV and mouse tissue extraction discussions), J. Ferreira da Silva and L. Hille (data analysis), L. Ma (cloning), R. Silverstein (coding and data analysis), H. Stutzman (gRNA cloning), and E. Eichelberger, R. Spellman, and W. das Neves (cell culture). We acknowledge M. Mabuchi and G.B. Robb for providing purified SpRY and SpRY-HF1 protein (generated as described previously^80^). This work was supported by a Charles A. King Trust Postdoctoral Research Fellowship, Bank of America, N.A., Co-Trustees (C.R.R.A.), a James L. and Elisabeth C. Gamble Endowed Fund for Neuroscience Research / Mass General Neuroscience Transformative Scholar Award (C.R.R.A.), a Massachusetts General Hospital (MGH) Executive Committee on Research (ECOR) Fund for Medical Discovery Fundamental Research Fellowship Award (K.A.C.), a Frederick Banting and Charles Best CIHR Doctoral Research Award (A.R.), a St. Jude Children’s Research Hospital Collaborative Research Consortium on Novel Gene Therapies for Sickle Cell Disease (S.Q.T.), a Muscular Dystrophy Association grant 575466 (R.K.), a Muscular Dystrophy Canada grant (R.K.), a Canadian Institutes of Health Research (CIHR) grant PJT-156379 (R.K.), an MGH Innovation Discovery Grant (to C.R.R.A. and B.P.K.), an MGH ECOR Howard M. Goodman Fellowship (B.P.K.), and National Institutes of Health (NIH) grants R01DC017117 (C.A.M.), U01AI157189 (S.Q.T.), P01HL142494 (B.P.K.) and DP2CA281401 (B.P.K.).

## Author contributions

C.R.R.A. and B.P.K. conceived of and designed the study. All authors designed, performed, or supervised experiments, and/or analyzed data. C.R.R.A., L.L.H., K.A.C., and R.M.D. performed cell culture, molecular, and biochemical experiments. D.L.C and C.A.M. produced AAVs. R.Y., A.B., A.R. and R.K. conducted the *in vivo* experiments with mice and analyzed data. C.R.L. and S.Q.T. designed and performed CHANGE-seq experiments. K.J.S. collected primary human fibroblasts and clinical data. C.R.R.A. and B.P.K. wrote the manuscript with contributions and/or revisions from all authors.

## Data availability

Primary datasets are available in **Supplementary Tables 1, 2, 6, and 7**. Sequencing datasets will be deposited with the NCBI Sequence Read Archive (SRA).

Some aspects of schematics in **Figures 4a** and **4d**, and **Supplementary Figure 17a**, were adapted from vector art provided by Servier Medical Art by Servier, which is licensed under a Creative Commons Attribution 3.0 Unported License (https://creativecommons.org/licenses/by/3.0/).”

## Methods

### Plasmids and oligonucleotides

Target site sequences for sgRNAs are available in **Supplementary Table 3**. Plasmids used in this study are described in **Supplementary Table 4**; new plasmids generated during this study have been deposited with Addgene (https://www.addgene.org/Benjamin_Kleinstiver/). Oligonucleotide sequences are available in **Supplementary Table 5**. Various ABE-SpRY plasmids were generated by subcloning the ABE8e or ABE8.20m^35,36^ deaminase sequence (Twist Biosciences) into the NotI and BglII sites of pCMV-T7-ABEmax(7.10)-SpRY-P2A-EGFP (RTW5025; Addgene plasmid 140003) via isothermal assembly^81^. We also similarly generated ABE8e and ABE8.20 versions of wild-type SpCas9, SpG, and SpCas9-NRRH, SpCas9-NRTH and SpCas9-NRCH, as well as ABE8e versions of SpCas9 and SpRY bearing HF1 mutations (N497A/R661A/Q695A/Q926A) or the HiFi mutation (R691A)^37,43-45^. Expression plasmids for human U6 promoter-driven sgRNAs were generated by annealing and ligating duplexed oligonucleotides corresponding to spacer sequences into BsmBI-digested pUC19-U6-BsmBI_cassette-SpCas9_sgRNA (BPK1520; Addgene plasmid 65777). Various Npu intein-split ABE constructs were cloned into N- and C-terminal AAV vectors (Addgene plasmids 137177 and 137178, respectively). The N-terminal vector was modified to include the ABE8e domain and additionally include the A61R mutation for SpRY. The C-terminal vector was modified to encode gRNAs A7, A8, and A10 and optionally to include the remainder of the SpRY mutations.

### Cell culture and transfections

Human HEK 293T cells (American Type Culture Collection; ATCC) were cultured in Dulbecco’s Modified Eagle Medium (DMEM) supplemented with 10% heat-inactivated FBS (HI-FBS) and 1% penicillin-streptomycin. Samples of supernatant media from cell culture experiments were analyzed monthly for the presence of mycoplasma using MycoAlert PLUS (Lonza).

For HEK 293T human cell experiments, transfections were performed 20 hours following seeding of 2×10^4^ HEK 293T cells per well in 96-well plates. Transfections containing 70 ng of ABE expression plasmid and 30 ng sgRNA expression plasmid mixed with 0.72 μL of TransIT-X2 (Mirus) in a total volume of 15 μL Opti-MEM (Thermo Fisher Scientific), incubated for 15 minutes at room temperature, and distributed across the seeded HEK 293T cells. Experiments were halted after 72 hours and genomic DNA (gDNA) was collected by discarding the media, resuspending the cells in 100 μL of quick lysis buffer (20 mM Hepes pH 7.5, 100 mM KCl, 5 mM MgCl_2_, 5% glycerol, 25 mM DTT, 0.1% Triton X-100, and 60 ng/μL Proteinase K (New England Biolabs; NEB)), heating the lysate for 6 minutes at 65 ºC, heating at 98 ºC for 2 minutes, and then storing at -20 ºC.

Fibroblasts were derived from skin biopsies from five different SMA patients. SMA type, SMN2 copy number and age at skin biopsy are provided in **Fig. 2a**. Fibroblasts were obtained from the Massachusetts General Hospital (MGH) SPOT SMA Longitudinal Population Database Repository (LPDR) database. Unique MGH IDs for fibroblasts lines 1 to 5 are #480, 570, 579, 603 and 571, respectively. Fibroblasts were cultured in Dulbecco’s Modified Eagle Medium (DMEM) supplemented with 10% HI-FBS and 1% penicillin/streptomycin. For experiments involving fluorescence-activated cell sorting, the media was modified to contain 20% HI-FBS for recovery after sorting. Fibroblasts were transfected with Lipofectamine LTX (ThermoFisher) to deliver separate plasmids encoding ABE8e-SpRY-P2A-EGFP and gRNA A8. Transfections were performed with a non-targeting gRNA to establish a “control” line and naïve cells were untreated. Approximately 48 hours after transfection, GFP+ fibroblasts were sorted (MGB HSCI CRM Flow Cytometry Core; BD FACS AriaIII cell sorter) and seeded into a pooled GFP+ population to grow for an additional 7 days. Two additional passages were performed to expand the sorted cells, which were then used to extract gDNA as described above at passages 3, 4, 5, and 6. In addition, extractions at passages 4, 5, and 6 were performed to extract RNA using the RNeasy Plus Universal Kit (Qiagen, Hilden, Germany) and protein using an SMA enzyme-linked immunosorbent assay (ELISA; Life Sciences Inc., Farmingdale, NY; ADI-900-209).

### Assessment of ABE activities in human cells

The efficiency of genome modification by ABEs were determined by next-generation sequencing using a 2-step PCR-based Illumina library construction method, similar to as previously described^37^. Briefly, genomic loci were amplified using approximately 50-100 ng of gDNA, Q5 High-fidelity DNA Polymerase (NEB), and PCR-1 primers (**Supplementary Table 5**) with cycling conditions of 1 cycle at 98 ºC for 2 min; 35 cycles of 98 ºC for 10 sec, 58 ºC for 10 sec, 72 ºC for 20 sec; and 1 cycle of 72 ºC for 1 min. PCR products were purified using paramagnetic beads prepared as previously described^82,83^. Approximately 20 ng of purified PCR-1 products were used as template for a second round of PCR (PCR-2) to add barcodes and Illumina adapter sequences using Q5 and primers (**Supplementary Table 5**) and cycling conditions of 1 cycle at 98 ºC for 2 min; 10 cycles at 98 ºC for 10 sec, 65 ºC for 30 sec, 72 ºC 30 sec; and 1 cycle at 72 ºC for 5 min. PCR products were purified prior to quantification via capillary electrophoresis (Qiagen QIAxcel), normalization, and pooling. Final libraries were quantified by qPCR using the KAPA Library Quantification Kit (Complete kit; Universal) (Roche) and sequenced on a MiSeq sequencer using a 300-cycle v2 kit (Illumina).

On-target genome editing activities were determined from sequencing data using CRISPResso2 (ref. ^84^) in pooled mode with custom input parameters for ABEs: -min_reads_to_use_region 100 -- quantification_window_size 10 --quantification_window_center -10 --base_editor_output -- min_frequency_alleles_around_cut_to_plot 0.05. Since amplification of *SMN2* also amplifies *SMN1*, final levels of editing were calculated as: ([%G in edited samples] - [%G in control samples]) / [%A in control samples] (**Supplementary Table 6**).

### Assessment of *SMN* transcript levels

RNA was extracted from fibroblasts using the RNeasy Plus Universal Kit (Qiagen, Hilden, Germany). RNA was reverse transcribed using the RT2 First Strand Kit (Qiagen, Hilden, Germany). For ddPCR reactions, cDNA was normalized to 2 ng/μL and each ddPCR reaction contained 12 ng of cDNA, 250 nM of each primer and 900 nM probe (**Supplementary Table 5**), and ddPCR supermix for probes (no dUTP) (BioRad). Droplets were generated using a QX200 Automated Droplet Generator (BioRad). Thermal cycling conditions were: 1 cycle at 95 °C for 10 min; 40 cycles at 94 °C for 30 sec, 58 °C for 1 min; and 1 cycle at 98 °C for 10 min; and hold at 4 °C. PCR products were analyzed using a QX200 Droplet Reader (BioRad) and absolute concentration was determined using QuantaSoft (v1.7.4). SMN exon 7 expression was calculated relative to the housekeeping gene (GAPDH) or total SMN transcript levels (exon 1-2 junction expression).

### Assessment of SMN protein levels

SMN protein levels were measured using an SMN-specific ELISA (Life Sciences Inc., Farmingdale, NY; ADI-900-209). Sample buffer provided with the ELISA kit was used to extract protein. In addition, SMN, PTEN and GAPDH protein levels were determined by immunoblotting as previously described^17^ with few modifications. For immunoblotting, cells were lysed using the buffer provided in the ELISA kit. In each well, 20 μg of protein was loaded in a 4-20% precast protein gel (Biorad, #4561096) and subjected to electrophoresis. Proteins were transferred to a PVDF membrane and blocked for 1 h at room temperature in Odyssey blocking buffer (Li-Cor, Lincoln, NE). Membranes were incubated with primary antibodies overnight at 4 °C. Primary antibodies were used to probe for SMN (BD Biosciences; 610647), PTEN (Cell Signaling; #9552) and GAPDH (Cell Signaling; #2118). Membranes were imaged using a ChemiDoc Touch System (Bio-Rad, USA) or the LI-COR Odyssey Infrared Imaging System (LI-COR, Inc., USA). SMN and PTEN expression were normalized by GAPDH expression.

### Off-target analysis

Circularization for high-throughput analysis of nuclease genome-wide effects by sequencing (CHANGE-seq) was performed as previously described^57^. Briefly, CHANGE-seq library preparation was performed on wild-type gDNA extracted from each of the five SMA-fibroblast lines (used for the on-target analysis) using the Gentra PureGene Tissue Kit (Qiagen). Approximately 5 μg of purified gDNA per CHANGE-seq reaction was tagmented with a custom Tn5-transposome^57^ to an average length of 400 bp, gap repaired with KAPA HiFi HotStart Uracil+ Ready Mix (Roche) and treated with a mixture of USER enzyme and T4 polynucleotide kinase (NEB). DNA was circularized at a concentration of 5 ng/μL with T4 DNA ligase (NEB), and treated with a cocktail of exonucleases, Lambda exonuclease (NEB), Exonuclease I (NEB) and Plasmid-Safe ATP-dependent DNase (Lucigen) to enzymatically degrade remaining linear DNA molecules. sgRNAs (Synthego) A7, A8, and A10 (**Supplementary Table 3**) were re-folded prior complexation with SpCas9 (Cas9 Nuclease, *S. pyogenes*; NEB), SpRY, or SpRY-HF1 (the latter two purified as previously described^80^) at a nuclease:sgRNA ratio of 1:3 to ensure full ribonucleoprotein complexation. *In vitro* cleavage reactions were performed in a 50 μL volume with NEB Buffer 3.1, 90 nM SpCas9, SpRY, or SpRY-HF1 protein, 270 nM of synthetic sgRNA and 125 ng of exonuclease treated circularized DNA. Digested products were treated with proteinase K (NEB), A-tailed (using KAPA High Throughput Library Preparation Kit; Roche), ligated with a hairpin adapter (NEB), treated with USER enzyme (NEB) and amplified by PCR using KAPA HiFi HotStart ReadyMix (Roche). Completed libraries were quantified by qPCR using KAPA Library Quantification kit (Complete kit; Universal) (Roche) and sequenced with 150 bp paired-end reads on an Illumina NextSeq 550 instrument. Data analysis was conducted as previously described^57^ (**Supplementary Table 2**).

For validation of potential off-target A-to-G editing, we selected the top 34 off-target sites with the highest average normalized CHANGE-seq read counts across the SpRY-A8 and SpRY-HF1-A8 treatments of the five SMA-fibroblasts experiments. Primers were designed for each target site using Primer3 (ref. ^85^) or were designed manually using genome.ucsc.edu and Geneious (v2021.2.2) for amplicons that failed to be designed by Primer3. For validation of potential off-target editing, gDNA was used as template for PCRs from three fibroblast cell lines (IDs 1, 2, and 3) that were either untreated or edited with ABE8e-SpRY and gRNA A8, and gDNA from untreated HEK 293Ts or those edited with ABE8e-SpRY and gRNA A8 or ABE8e-SpRY-HF1 and gRNA A8. Off-target sites were amplified from ∼100 ng gDNA using primers (**Supplementary Table 5**) and the PCR-1 parameters described above with modified cycling conditions where appropriate (**Supplementary Table 5**). Following PCR-1, amplicons were cleaned up, quantified, and subject to library construction using the KAPA HyperPrep Kit (Roche) with Illumina-competent adapters (**Supplementary Table 5**. Amplicons were sequenced and data analyzed as described above for assessment of ABE activities in human cells. Statistical analyses were conducted using Graph Pad Prism 8 (Graph Pad Software Inc.) (**Supplementary Table 2**). Multiple t-tests were used to compare groups. Statistical significance was set at p < 0.05.

### AAV production

For tissue transduction experiments, AAV9 and AAV-F vectors expressing eGFP were produced in-house. The AAV9 capsid was encoded in the pAR9 rep/cap vector kindly provided by Dr. Miguel Sena-Esteves at the University of Massachusetts Medical School (Worcester, MA). AAV-F^60,86^ is an engineered AAV9-based capsid in pAR9 (rep/cap) (Addgene plasmid 166921). AAV production was performed as previously described^60^. Briefly, HEK 293T cells were transfected using polyethylenimine and three plasmids encoding the AAV-rep/cap for either AAV9 or AAV-F, an adenovirus helper plasmid (pAdΔF6; Addgene plasmid 112867), and an ITR-flanked AAV transgene expression plasmid AAV-CAG-eGFP (provided by Dr. Miguel Sena-Esteves). The latter plasmid contains AAV inverted terminal repeats (ITRs) flanking the CAG expression cassette which consists of a hybrid CMV-IE enhancer, chicken β-actin (CBA) promoter, a beta actin exon, chimeric intron, eGFP cDNA, a woodchuck hepatitis virus post-translational response element (WPRE) and tandem SV40 and bGH poly-A signal sequences. Cell lysates as well as polyethylene glycol (PEG) precipitated vector from culture media were harvested 68-72 hr post transfection and purified by ultracentrifugation of an iodixanol density gradient. Iodixanol was removed and buffer exchanged to phosphate buffered saline (PBS) containing 0.001% Pluronic F68 (Gibco) using 7 kDa molecular weight cutoff Zeba desalting columns (ThermoFisher Scientific). AAV was concentrated using Amicon Ultra-2 100 kDa MWCO ultrafiltration devices (Millipore Sigma). Vector titers in vg/mL were determined by qPCR using an ABI Fast 7500 Real-time PCR system (Applied Biosystems), with Applied Biosystems TaqMan Fast Universal PCR Master Mix 2x, No AmpErase UNG (ThermoFisher Scientific) using primers and a probe to the bovine growth hormone (bGH) polyadenylation signal sequence (**Supplementary Table 5**), with cycling parameters of 95 °C for 20 s followed by 40 cycles of 95 °C for 3 s and 60 °C for 30 s. Titers were interpolated from a standard curve made with a XbaI-linearized AAV-CAG-eGFP plasmid. Vectors were pipetted into single-use aliquots and stored at -80°C.

For genome editing experiments, two AAV9 vectors encoding ABE8e-SpRY split into N-term and C-terminal fragments via an Npu intein (as described above and similar to as previously reported^59^) paired with gRNA A8 were packaged by PackGene Biotech Inc. (Worcester, MA).

### *In vivo* experiments in mice

SMNΔ7 mice (FVB.Cg-Grm7^Tg(SMN2)89Ahmb^ Smn1^tm1Msd^ Tg (SMN2*delta7) 4299Ahmb/J) were housed and bred at the University of Ottawa Animal Care Facility. Mice were housed in a 12h/12h light/dark cycle with access to food and water *ad libitum*. This study was approved by the Animal Care and Veterinary Services of the University of Ottawa, ON, Canada and all animals were cared for according to the Canadian Council on Animal Care.

For tissue transduction tests, intracerebroventricular (ICV) injections were performed in two litters of SMNΔ7 mice at postnatal day 1 (P1). For each mouse, 3 μL ICV injections contained 2 × 10^10^ vg AAV9-EGFP or AAV-F-EGFP. Mice were sacrificed at P13. Brain and spinal cord were collected, fixed in 4% paraformaldehyde (overnight at 4 °C), transferred to 30% sucrose (overnight at 4 °C), embedded in OCT, and frozen in liquid nitrogen. Cryosections of 16 μm were mounted on slides and kept at -20°C until staining, when the slides were air dried at room temperature (RT) for 24 h and rinsed with PBS for 3 × 5 min. Samples were permeabilized in 0.5% Triton X-100 in PBS for 25 min, then incubated in blocking solution (1% BSA, 10% goat serum, and 0.2% Triton X-100) for 40 min at room temperature (RT). Sections were then incubated with anti-GFP antibody at a dilution of 1:1,000 (Thermo Fisher Scientific, #A11122) in blocking solution overnight at 4 °C. After the first antibody incubation, slides were washed 3 × 10 min with PBS at RT. Samples were then incubated with Alexa Fluor goat anti-rabbit 488 (Thermo Fisher Scientific, ***#***A-11008) at a dilution of 1:100 for 1 h at RT. Nuclei were counterstained with 40,6-diamidino-2-phenylindole (DAPI) 1:1,000 in TBST for 5 min. Finally, slides were carefully rinsed 3 × 5 min with PBS and slides were mounted with Fluorescent Mounting Medium (Dako) and examined under fluorescence using a Zeiss microscope equipped with an AxioCam HRm camera.

For western blot analysis, tissue processing and immunoblotting were performed as previously described^87^. After euthanasia at P13, brain, spinal cord, liver, and heart were dissected, and flash frozen in liquid nitrogen. Protein was extracted from frozen tissue by homogenization of tissue with RIPA lysis buffer and PMSF (Cell Signaling, Danvers, MA). Protein concentrations of samples were determined using Bradford Assay (Bio-Rad, Hercules, CA). 20 μg of protein was loaded onto a 12% acrylamide gel and subjected to sodium dodecyl sulfate polyacrylamide gel electrophoresis. Proteins were transferred to a PVDF membrane (Immobilon-FL, Millipore, Burlington, MA) and blocked for 1 h at room temperature in Odyssey blocking buffer (Li-Cor, Lincoln, NE). Blots were incubated with anti-GFP (1:2,000; A11122 ThermoFisher Scientific). Signals were detected with Odyssey CLx (Li-Cor). Raw values were normalized by geometric mean and used for subsequent housekeeping normalization (α-tubulin values; 1:10,000 mouse anti-tubulin; Calbiochem CP06) from the same blot.

For genome editing experiments, 3 μL ICV injections were performed in a litter of SMNΔ7 mice at P1 with a dose of 3 × 10^10^ vg total N- and C-terminal AAV9 constructs. A control litter of SMNΔ7 mice were left uninjected. Mice were weighed every 2 days and sacrificed at P13 for collection of various tissues including brain, spinal cord, liver, heart, and skeletal muscle. DNA was extracted from each tissue using the Agencourt DNAdvance kit (Beckman Coulter, CA). On-target editing in the tissues was analyzed from extracted gDNA by amplifying the human SMN sequence as described above for assessment of ABE activities in human cells.

## References

1 Groen, E. J. N., Talbot, K. & Gillingwater, T. H. Advances in therapy for spinal muscular atrophy: promises and challenges. Nat Rev Neurol 14, 214–224, doi:10.1038/nrneurol.2018.4 (2018).

2 Bergin, A. et al. Identification and characterization of a mouse homologue of the spinal muscular atrophy-determining gene, survival motor neuron. Gene 204, 47–53, doi:10.1016/s0378-1119(97)00510-6 (1997).

3 Ogino, S. & Wilson, R. B. Genetic testing and risk assessment for spinal muscular atrophy (SMA). Hum Genet 111, 477–500, doi:10.1007/s00439-002-0828-x (2002).

4 Crawford, T. O. et al. Evaluation of SMN protein, transcript, and copy number in the biomarkers for spinal muscular atrophy (BforSMA) clinical study. PLoS One 7, e33572, doi:10.1371/journal.pone.0033572 (2012).

5 Mailman, M. D. et al. Molecular analysis of spinal muscular atrophy and modification of the phenotype by SMN2. Genet Med 4, 20–26, doi:10.1097/00125817-200201000-00004 (2002).

6 Feldkotter, M., Schwarzer, V., Wirth, R., Wienker, T. F. & Wirth, B. Quantitative analyses of SMN1 and SMN2 based on real-time lightCycler PCR: fast and highly reliable carrier testing and prediction of severity of spinal muscular atrophy. Am J Hum Genet 70, 358–368, doi:10.1086/338627 (2002).

7 Finkel, R. S. et al. Treatment of infantile-onset spinal muscular atrophy with nusinersen: a phase 2, open-label, dose-escalation study. Lancet 388, 3017–3026, doi:10.1016/S0140-6736(16)31408-8 (2016).

8 Finkel, R. S. et al. Nusinersen versus Sham Control in Infantile-Onset Spinal Muscular Atrophy. N Engl J Med 377, 1723–1732, doi:10.1056/NEJMoa1702752 (2017).

9 Mercuri, E. et al. Nusinersen versus Sham Control in Later-Onset Spinal Muscular Atrophy. N Engl J Med 378, 625–635, doi:10.1056/NEJMoa1710504 (2018).

10 Sturm, S. et al. A phase 1 healthy male volunteer single escalating dose study of the pharmacokinetics and pharmacodynamics of risdiplam (RG7916, RO7034067), a SMN2 splicing modifier. Br J Clin Pharmacol 85, 181–193, doi:10.1111/bcp.13786 (2019).

11 Darras, B. T. et al. Risdiplam-Treated Infants with Type 1 Spinal Muscular Atrophy versus Historical Controls. N Engl J Med 385, 427–435, doi:10.1056/NEJMoa2102047 (2021).

12 Baranello, G. et al. Risdiplam in Type 1 Spinal Muscular Atrophy. N Engl J Med 384, 915–923, doi:10.1056/NEJMoa2009965 (2021).

13 Ratni, H., Scalco, R. S. & Stephan, A. H. Risdiplam, the First Approved Small Molecule Splicing Modifier Drug as a Blueprint for Future Transformative Medicines. ACS Med Chem Lett 12, 874–877, doi:10.1021/acsmedchemlett.0c00659 (2021).

14 De Vivo, D. C. et al. Nusinersen initiated in infants during the presymptomatic stage of spinal muscular atrophy: Interim efficacy and safety results from the Phase 2 NURTURE study. Neuromuscul Disord 29, 842–856, doi:10.1016/j.nmd.2019.09.007 (2019).

15 Hua, Y. et al. Peripheral SMN restoration is essential for long-term rescue of a severe spinal muscular atrophy mouse model. Nature 478, 123–126, doi:10.1038/nature10485 (2011).

16 Lipnick, S. L. et al. Systemic nature of spinal muscular atrophy revealed by studying insurance claims. PLoS One 14, e0213680, doi:10.1371/journal.pone.0213680 (2019).

17 Nery, F. C. et al. Impaired kidney structure and function in spinal muscular atrophy. Neurol Genet 5, e353, doi:10.1212/NXG.0000000000000353 (2019).

18 Kim, J. K. et al. Muscle-specific SMN reduction reveals motor neuron-independent disease in spinal muscular atrophy models. J Clin Invest 130, 1271–1287, doi:10.1172/JCI131989 (2020).

19 Mendell, J. R. et al. Single-Dose Gene-Replacement Therapy for Spinal Muscular Atrophy. N Engl J Med 377, 1713–1722, doi:10.1056/NEJMoa1706198 (2017).

20 Thomsen, G. et al. Biodistribution of onasemnogene abeparvovec DNA, mRNA and SMN protein in human tissue. Nat Med 27, 1701–1711, doi:10.1038/s41591-021-01483-7 (2021).

21 Alves, C. R. R. et al. Whole blood survival motor neuron protein levels correlate with severity of denervation in spinal muscular atrophy. Muscle Nerve 62, 351–357, doi:10.1002/mus.26995 (2020).

22 Van Alstyne, M. et al. Gain of toxic function by long-term AAV9-mediated SMN overexpression in the sensorimotor circuit. Nat Neurosci, doi:10.1038/s41593-021-00827-3 (2021).

23 Zhou, M. et al. Seamless Genetic Conversion of SMN2 to SMN1 via CRISPR/Cpf1 and Single-Stranded Oligodeoxynucleotides in Spinal Muscular Atrophy Patient-Specific Induced Pluripotent Stem Cells. Hum Gene Ther 29, 1252–1263, doi:10.1089/hum.2017.255 (2018).

24 Li, J. J. et al. Disruption of splicing-regulatory elements using CRISPR/Cas9 to rescue spinal muscular atrophy in human iPSCs and mice. Natl Sci Rev 7, 92–101, doi:10.1093/nsr/nwz131 (2020).

25 Miccio, A., Antoniou, P., Ciura, S. & Kabashi, E. Novel genome-editing-based approaches to treat motor neuron diseases: Promises and challenges. Mol Ther 30, 47–53, doi:10.1016/j.ymthe.2021.04.003 (2022).

26 Singh, N. N., Howell, M. D., Androphy, E. J. & Singh, R. N. How the discovery of ISS-N1 led to the first medical therapy for spinal muscular atrophy. Gene Ther 24, 520–526, doi:10.1038/gt.2017.34 (2017).

27 Kosicki, M., Tomberg, K. & Bradley, A. Repair of double-strand breaks induced by CRISPR-Cas9 leads to large deletions and complex rearrangements. Nat Biotechnol 36, 765–771, doi:10.1038/nbt.4192 (2018).

28 Leibowitz, M. L. et al. Chromothripsis as an on-target consequence of CRISPR-Cas9 genome editing. Nat Genet 53, 895–905, doi:10.1038/s41588-021-00838-7 (2021).

29 Alanis-Lobato, G. et al. Frequent loss of heterozygosity in CRISPR-Cas9-edited early human embryos. Proc Natl Acad Sci U S A 118, doi:10.1073/pnas.2004832117 (2021).

30 Enache, O. M. et al. Cas9 activates the p53 pathway and selects for p53-inactivating mutations. Nat Genet 52, 662–668, doi:10.1038/s41588-020-0623-4 (2020).

31 Morgens, D. W. et al. Genome-scale measurement of off-target activity using Cas9 toxicity in high-throughput screens. Nat Commun 8, 15178, doi:10.1038/ncomms15178 (2017).

32 Anzalone, A. V., Koblan, L. W. & Liu, D. R. Genome editing with CRISPR-Cas nucleases, base editors, transposases and prime editors. Nat Biotechnol 38, 824–844, doi:10.1038/s41587-020-0561-9 (2020).

33 Gaudelli, N. M. et al. Programmable base editing of A*T to G*C in genomic DNA without DNA cleavage. Nature 551, 464–471, doi:10.1038/nature24644 (2017).

34 Komor, A. C., Kim, Y. B., Packer, M. S., Zuris, J. A. & Liu, D. R. Programmable editing of a target base in genomic DNA without double-stranded DNA cleavage. Nature 533, 420–424, doi:10.1038/nature17946 (2016).

35 Gaudelli, N. M. et al. Directed evolution of adenine base editors with increased activity and therapeutic application. Nat Biotechnol 38, 892–900, doi:10.1038/s41587-020-0491-6 (2020).

36 Richter, M. F. et al. Phage-assisted evolution of an adenine base editor with improved Cas domain compatibility and activity. Nat Biotechnol 38, 883–891, doi:10.1038/s41587-020-0453-z (2020).

37 Walton, R. T., Christie, K. A., Whittaker, M. N. & Kleinstiver, B. P. Unconstrained genome targeting with near-PAMless engineered CRISPR-Cas9 variants. Science 368, 290–296, doi:10.1126/science.aba8853 (2020).

38 Koblan, L. W. et al. Improving cytidine and adenine base editors by expression optimization and ancestral reconstruction. Nat Biotechnol 36, 843–846, doi:10.1038/nbt.4172 (2018).

39 Ma, H. et al. Pol III Promoters to Express Small RNAs: Delineation of Transcription Initiation. Mol Ther Nucleic Acids 3, e161, doi:10.1038/mtna.2014.12 (2014).

40 Gao, Z., Harwig, A., Berkhout, B. & Herrera-Carrillo, E. Mutation of nucleotides around the +1 position of type 3 polymerase III promoters: The effect on transcriptional activity and start site usage. Transcription 8, 275–287, doi:10.1080/21541264.2017.1322170 (2017).

41 Kim, S., Bae, T., Hwang, J. & Kim, J. S. Rescue of high-specificity Cas9 variants using sgRNAs with matched 5’ nucleotides. Genome Biol 18, 218, doi:10.1186/s13059-017-1355-3 (2017).

42 Arbab, M. et al. Determinants of Base Editing Outcomes from Target Library Analysis and Machine Learning. Cell 182, 463–480 e430, doi:10.1016/j.cell.2020.05.037 (2020).

43 Miller, S. M. et al. Continuous evolution of SpCas9 variants compatible with non-G PAMs. Nat Biotechnol 38, 471–481, doi:10.1038/s41587-020-0412-8 (2020).

44 Vakulskas, C. A. et al. A high-fidelity Cas9 mutant delivered as a ribonucleoprotein complex enables efficient gene editing in human hematopoietic stem and progenitor cells. Nat Med 24, 1216–1224, doi:10.1038/s41591-018-0137-0 (2018).

45 Kleinstiver, B. P. et al. High-fidelity CRISPR-Cas9 nucleases with no detectable genome-wide off-target effects. Nature 529, 490–495, doi:10.1038/nature16526 (2016).

46 Kashima, T., Rao, N. & Manley, J. L. An intronic element contributes to splicing repression in spinal muscular atrophy. Proc Natl Acad Sci U S A 104, 3426–3431, doi:10.1073/pnas.0700343104 (2007).

47 Singh, N. K., Singh, N. N., Androphy, E. J. & Singh, R. N. Splicing of a critical exon of human Survival Motor Neuron is regulated by a unique silencer element located in the last intron. Mol Cell Biol 26, 1333–1346, doi:10.1128/MCB.26.4.1333-1346.2006 (2006).

48 Cartegni, L., Hastings, M. L., Calarco, J. A., de Stanchina, E. & Krainer, A. R. Determinants of exon 7 splicing in the spinal muscular atrophy genes, SMN1 and SMN2. Am J Hum Genet 78, 63–77, doi:10.1086/498853 (2006).

49 Kashima, T., Rao, N., David, C. J. & Manley, J. L. hnRNP A1 functions with specificity in repression of SMN2 exon 7 splicing. Hum Mol Genet 16, 3149–3159, doi:10.1093/hmg/ddm276 (2007).

50 Kashima, T. & Manley, J. L. A negative element in SMN2 exon 7 inhibits splicing in spinal muscular atrophy. Nat Genet 34, 460–463, doi:10.1038/ng1207 (2003).

51 Godena, V. K. & Ning, K. Phosphatase and tensin homologue: a therapeutic target for SMA. Signal Transduct Target Ther 2, 17038, doi:10.1038/sigtrans.2017.38 (2017).

52 Little, D. et al. PTEN depletion decreases disease severity and modestly prolongs survival in a mouse model of spinal muscular atrophy. Mol Ther 23, 270–277, doi:10.1038/mt.2014.209 (2015).

53 Bae, S., Park, J. & Kim, J. S. Cas-OFFinder: a fast and versatile algorithm that searches for potential off-target sites of Cas9 RNA-guided endonucleases. Bioinformatics 30, 1473–1475, doi:10.1093/bioinformatics/btu048 (2014).

54 Jinek, M. et al. A programmable dual-RNA-guided DNA endonuclease in adaptive bacterial immunity. Science 337, 816–821, doi:10.1126/science.1225829 (2012).

55 Jiang, W., Bikard, D., Cox, D., Zhang, F. & Marraffini, L. A. RNA-guided editing of bacterial genomes using CRISPR-Cas systems. Nat Biotechnol 31, 233–239, doi:10.1038/nbt.2508 (2013).

56 Kleinstiver, B. P. et al. Engineered CRISPR-Cas9 nucleases with altered PAM specificities. Nature 523, 481–485, doi:10.1038/nature14592 (2015).

57 Lazzarotto, C. R. et al. CHANGE-seq reveals genetic and epigenetic effects on CRISPR-Cas9 genome-wide activity. Nat Biotechnol 38, 1317–1327, doi:10.1038/s41587-020-0555-7 (2020).

58 Koblan, L. W. et al. In vivo base editing rescues Hutchinson-Gilford progeria syndrome in mice. Nature 589, 608–614, doi:10.1038/s41586-020-03086-7 (2021).

59 Levy, J. M. et al. Cytosine and adenine base editing of the brain, liver, retina, heart and skeletal muscle of mice via adeno-associated viruses. Nat Biomed Eng 4, 97–110, doi:10.1038/s41551-019-0501-5 (2020).

60 Hanlon, K. S. et al. Selection of an Efficient AAV Vector for Robust CNS Transgene Expression. Mol Ther Methods Clin Dev 15, 320–332, doi:10.1016/j.omtm.2019.10.007 (2019).

61 Le, T. T. et al. SMNDelta7, the major product of the centromeric survival motor neuron (SMN2) gene, extends survival in mice with spinal muscular atrophy and associates with full-length SMN. Hum Mol Genet 14, 845–857, doi:10.1093/hmg/ddi078 (2005).

62 Lutz, C. M. et al. Postsymptomatic restoration of SMN rescues the disease phenotype in a mouse model of severe spinal muscular atrophy. J Clin Invest 121, 3029–3041, doi:10.1172/JCI57291 (2011).

63 Kray, K. M., McGovern, V. L., Chugh, D., Arnold, W. D. & Burghes, A. H. M. Dual SMN inducing therapies can rescue survival and motor unit function in symptomatic Δ7SMA mice. Neurobiol Dis 159, 105488, doi:10.1016/j.nbd.2021.105488 (2021).

64 Banskota, S. et al. Engineered virus-like particles for efficient in vivo delivery of therapeutic proteins. Cell 185, 250–265 e216, doi:10.1016/j.cell.2021.12.021 (2022).

65 Buettner, J. M. et al. Central synaptopathy is the most conserved feature of motor circuit pathology across spinal muscular atrophy mouse models. iScience 24, 103376, doi:10.1016/j.isci.2021.103376 (2021).

66 Thomas, N. H. & Dubowitz, V. The natural history of type I (severe) spinal muscular atrophy. Neuromuscul Disord 4, 497–502, doi:10.1016/0960-8966(94)90090-6 (1994).

67 Swoboda, K. J. et al. Natural history of denervation in SMA: relation to age, SMN2 copy number, and function. Ann Neurol 57, 704–712, doi:10.1002/ana.20473 (2005).

68 Frangoul, H. et al. CRISPR-Cas9 Gene Editing for Sickle Cell Disease and beta-Thalassemia. N Engl J Med 384, 252–260, doi:10.1056/NEJMoa2031054 (2021).

69 Gillmore, J. D. et al. CRISPR-Cas9 In Vivo Gene Editing for Transthyretin Amyloidosis. N Engl J Med 385, 493–502, doi:10.1056/NEJMoa2107454 (2021).

70 Gao, Y. et al. Systematic characterization of short intronic splicing-regulatory elements in SMN2 pre-mRNA. Nucleic Acids Res 50, 731–749, doi:10.1093/nar/gkab1280 (2022).

71 Goertsen, D. et al. AAV capsid variants with brain-wide transgene expression and decreased liver targeting after intravenous delivery in mouse and marmoset. Nat Neurosci 25, 106–115, doi:10.1038/s41593-021-00969-4 (2022).

72 Ravindra Kumar, S. et al. Multiplexed Cre-dependent selection yields systemic AAVs for targeting distinct brain cell types. Nat Methods 17, 541–550, doi:10.1038/s41592-020-0799-7 (2020).

73 Stanton, A. C. et al. Systemic administration of novel engineered AAV capsids facilitates enhanced transgene expression in the macaque CNS. Med (N Y) 4, 31–50 e38, doi:10.1016/j.medj.2022.11.002 (2023).

74 Rossidis, A. C. et al. In utero CRISPR-mediated therapeutic editing of metabolic genes. Nat Med 24, 1513–1518, doi:10.1038/s41591-018-0184-6 (2018).

75 Poirier, A. et al. Risdiplam distributes and increases SMN protein in both the central nervous system and peripheral organs. Pharmacol Res Perspect 6, e00447, doi:10.1002/prp2.447 (2018).

76 Zhao, X. et al. Pharmacokinetics, pharmacodynamics, and efficacy of a small-molecule SMN2 splicing modifier in mouse models of spinal muscular atrophy. Hum Mol Genet 25, 1885–1899, doi:10.1093/hmg/ddw062 (2016).

77 Passini, M. A. et al. Antisense oligonucleotides delivered to the mouse CNS ameliorate symptoms of severe spinal muscular atrophy. Sci Transl Med 3, 72ra18, doi:10.1126/scitranslmed.3001777 (2011).

78 Chatterjee, P. et al. A Cas9 with PAM recognition for adenine dinucleotides. Nat Commun 11, 2474, doi:10.1038/s41467-020-16117-8 (2020).

79 Vicencio, J. et al. Genome editing in animals with minimal PAM CRISPR-Cas9 enzymes. Nat Commun 13, 2601, doi:10.1038/s41467-022-30228-4 (2022).

80 Christie, K. A. et al. Precise DNA cleavage using CRISPR-SpRYgests. Nat Biotechnol, doi:10.1038/s41587-022-01492-y (2022).

81 Gibson, D. G. et al. Enzymatic assembly of DNA molecules up to several hundred kilobases. Nat Methods 6, 343–345, doi:10.1038/nmeth.1318 (2009).

82 Kleinstiver, B. P. et al. Engineered CRISPR-Cas12a variants with increased activities and improved targeting ranges for gene, epigenetic and base editing. Nat Biotechnol 37, 276–282, doi:10.1038/s41587-018-0011-0 (2019).

83 Rohland, N. & Reich, D. Cost-effective, high-throughput DNA sequencing libraries for multiplexed target capture. Genome Res 22, 939–946, doi:10.1101/gr.128124.111 (2012).

84 Clement, K. et al. CRISPResso2 provides accurate and rapid genome editing sequence analysis. Nat Biotechnol 37, 224–226, doi:10.1038/s41587-019-0032-3 (2019).

85 Untergasser, A. et al. Primer3--new capabilities and interfaces. Nucleic Acids Res 40, e115, doi:10.1093/nar/gks596 (2012).

86 Beharry, A. et al. The AAV9 Variant Capsid AAV-F Mediates Widespread Transgene Expression in Nonhuman Primate Spinal Cord After Intrathecal Administration. Hum Gene Ther 33, 61–75, doi:10.1089/hum.2021.069 (2022).

87 Reilly, A. et al. Central and peripheral delivered AAV9-SMN are both efficient but target different pathomechanisms in a mouse model of spinal muscular atrophy. Gene Ther 29, 544–554, doi:10.1038/s41434-022-00338-1 (2022).

