## Supplementary Materials for "Base editing as a genetic treatment for spinal muscular atrophy"

Supplementary Tables

**Supplementary Table 1**: CHANGE-seq data summary

**Supplementary Table 2**: Off-target sequencing and results of statistical tests

**Supplementary Table 3**: gRNA target sites

**Supplementary Table 4**: Plasmids

**Supplementary Table 5**: Oligonucleotides and probes

**Supplementary Table 6**: Base editor data summary

**Supplementary Table 7**: Primary datasets

***Note:*** *All Supplementary Tables are attached as separate .xlsx files*.

Supplementary Notes

**Supplementary Note 1**: Analysis of the impact of 5' gRNA spacer architectures p.2

**Supplementary Note 2**: Assessment of ABEs on targets with adenine homopolymers p.2

**Supplementary Note 3**: Analysis of *SMN2* intron 7 ISS targeting p.3

**Supplementary Note 4**: CHANGE-seq data analysis p.3

**Supplementary Note 5**: Comparison of AAV9 and AAV-F capsids p.4

**Supplementary Note 6**: Details and results of *in vivo* SMNΔ7 mice injections p.5

Supplementary Figures and Legends

**Supplementary Figures 1-17** p.7

Supplementary References p.27

### Supplementary Notes:

#### Supplementary Note 1: Analysis of the impact of 5' gRNA spacer architectures.

To determine whether we could enhance on-target editing of C6T in *SMN2* exon 7, we examined different gRNA architectures that varied in the composition of the 5' end of the spacer. gRNA expression is generally regarded as most efficient from a U6 promoter when a guanine (G) base is present to initiate transcription[1,2]. Our previous experiments utilized gRNAs with 20 nucleotides spacers that, when necessary, harbored a mismatched 5' base that was obligately substituted to a G. An additional method to construct gRNAs with 5' Gs is to append an additional 5' G, generating gRNAs with 21 nucleotides spacers[3]. We compared the efficiency of *SMN2* base editing when using either 20 nucleotides spacers with 5' Gs or 21 nucleotides spacers with added Gs (+1 5'G). In general, we observed comparable on-target editing of C6T irrespective of the gRNA architecture for experiments using ABE8e-SpCas9 (**Sup. Fig. 1a**) and ABE8e-SpRY (**Sup. Fig. 1b**).

As a complementary analysis, we examined the impact of 5' gRNA architecture on *SMN2* C6T base editing when using conventional ABE8e-SpCas9 or ABE8e-SpRY constructs or their analogous HF1 and HiFi versions[4,5]. With the 20 nt 5'G spacers, we observed a loss in editing with ABE8e-SpCas9-HF1 or ABE8e-SpCas9-HiFi when using gRNA A10 (**Sup. Fig. 4a**). These results are consistent with a sensitivity of high-fidelity variants to PAM distal mismatches in the spacer, including mismatches required for gRNA transcription, consistent with prior reports[3,6,7]. It is also possible that high-fidelity base editors are more affected by mismatched 5' spacers when the target base is near the border of the ABE edit window[8]. In experiments using ABE8e-SpRY-HF1 or ABE8e-SpRY-HiFi and gRNAs A5, A7, or A8 harboring the 20 nt 5'G spacers, in most cases we observed comparable editing to ABE8e-SpRY (**Sup. Fig. 4b**). Next, when comparing the 20 nt 5'G and 21 nt +1 5'G spacer architectures, we observed low levels of editing with both ABE8e-SpCas9-HF1 or ABE8e-SpCas9-HiFi (**Sup. Figs. 4d** and **4d**, respectively). Experiments with ABE8e-SpRY-HF1 or ABE8e-SpRY-HiFi (**Sup. Figs. 4e** and **4f**, respectively) revealed similar levels of C6T editing irrespective of the 5' gRNA spacer architecture with SpRY high-fidelity variants.

#### Supplementary Note 2: Assessment of ABEs on targets with adenine homopolymers.

Our observation of minimal bystander editing of the three other adenines directly adjacent to the C6T edit can only be partially explained by the positioning of the additional adenines outside or at the border of the conventional ABE8e edit window[8]. Reduced bystander editing of adenines adjacent to and 3' of the primary adenine is partially supported by a previous machine learning model that indicated preceding adenines are modestly inhibitory to ABE efficiency[9]. We performed additional experiments to compare the A-to-G conversion efficiencies of the ABEmax[10,11], ABE8.20m[12], and ABE8e[8] deaminases fused to wild-type SpCas9 (**Sup. Figs. 2a** and **2b**), SpG[13] (**Sup. Figs. 2c** and **2d**) or SpRY[13] (**Sup. Figs. 2e** and **2f**), using gRNAs targeting sites with adenines interspersed with other nucleotides (**Sup. Figs. 2a**, **2c**, and **2e**) or gRNAs targeting sites containing multiple consecutive adenines (**Sup. Figs. 2b**, **2d**, and **2f**). In nearly all cases (with different gRNAs or SpCas9

PAM variants), we observed the highest levels of editing and widest edit windows with ABE8e followed by ABE8.20m, with both deaminases being substantially superior to ABEmax in terms of A-to-G editing efficiency. Across the sites with consecutive adenines (**Sup. Figs. 2b**, **2d**, and **2f**), we observed editing efficiencies mostly consistent with the edit windows of each deaminase on the sites with interspersed adenines (**Sup. Figs. 2a**, **2c**, and **2e**), and only minimal reduction of adenines that were flanked by 5' adenines.

**Supplementary Note 3: Analysis of *SMN2* intron 7 ISS targeting.**

In experiments targeting the ISS-N1 motif, we observed A-to-G editing of several adenines while using various gRNAs (**Sup. Fig. 5c**). The predominantly edited nucleotide was the first adenine in the ISS-N1 motif with >15% A-to-G editing by ABE8e-SpRY when using ISS-N1-gRNA1 (**Sup. Fig. 5c**). We also observed editing of several adenines within the ISS+100 motif, depending on the gRNA (**Sup. Fig. 5d**). When using ISS+100-gRNA4, treatment with ABE8e-SpRY led to greater than 40% editing in the fourth adenine of this motif (**Sup. Fig. 5d**).

Next, we explored multiplex editing via co-transfection of the ISS-targeted gRNAs along with the *SMN2* C6T targeted gRNA A8. We observed similar levels of adenine editing within the ISS-N1 (**Sup. Fig. 5c**) and ISS+100 (**Sup. Fig. 5c**) motifs when comparing single ISS gRNA targeting versus multiplex ISS-C6T targeting, with slightly reduced editing in the multiplex conditions, likely due to the use of half the effective dose of ISS gRNAs in transfections. For the multiplex edited samples, we observed robust editing at the C6T target adenine in *SMN2* when using gRNA A8 paired with gRNAs for either ISS-N1 or ISS+100 (**Sup. Figs. 5e** and **5f**, respectively), at levels comparable to transfections with the SMN2 C6T gRNA A8 alone. Thus, multiplex editing does not substantially reduce editing of *SMN2* C6T, suggesting that the A8 gRNA is less sensitive to gRNA dose.

**Supplementary Note 4: CHANGE-seq data analysis and validated off-target sites**

When performing CHANGE-seq experiments, the number of total reads returned from an experiment is a function of several factors including the total number of off-target sites, the efficiency of cleavage at the on- and off-target site(s), but also importantly, the experimental efficiency and sequencing depth of that individual experiment. Aside from the confounding variable of utilizing gRNAs targeting distinct sites, our observation of an increased number of total number of off-target sites with SpRY compared to WT SpCas9 (**Fig. 3b**) is consistent with the hypothesis that SpCas9 enzymes with minimal PAM requirements should encounter an expanded superset of off-target sites[13,14] (**Fig. 3a**). However, to some degree this might also reflect an increased sequencing efficiency of the SpRY-based experiments, given that the overall number of reads for some SpRY or SpRY-HF1 nucleases with gRNAs A7 and gRNA A8 samples (**Sup. Figs. 10-13**) are substantially higher than the number of reads for the WT SpCas9 nuclease with gRNA A10 samples (**Sup. Figs. 8** and **9**; see also summary graphs below). Thus, when analyzing the CHANGE-seq outputs from our experiments, one method to normalize the data is to focus on off-target sites that occur at a >1% abundance (of total CHANGE-seq reads for that experiment). This may avoid over-interpretation of lower efficiency sites that are near floor of detection (either due to low cleavage efficiency or from expanded sampling due to higher overall sequencing). Prior evidence suggests that the rank-order of CHANGE-seq detected sites is generally representative of editing levels in cells[15,16] making the calibration of focusing on the top ranked sites a reasonable method to analyze the off-target sites that are most likely to be edited (without bias due to sequencing depth of that run).

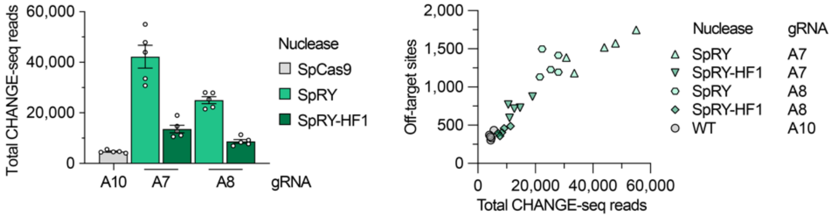


We observed one highly abundant CHANGE-seq off-target detected for WT SpCas9 nuclease and gRNA A10 that was located on chromosome 7 (31861836-31861859 using GRCh38) in an intron of *PPP1R17* (with an average of 10.78-fold greater reads at the off-target vs. on-target site, averaged across the five CHANGE-seq replicates; **Sup. Figs. 8** and **9**), With SpRY nuclease and gRNA A7, we observed one off-target site that was cleaved approximately as efficiently as the on-target that was located on chromosome 13 (94057979-94058002) in an intron of *GPC6* (1.04-fold difference; **Sup. Figs. 10** and **11**). Notably, this off-target was dramatically reduced by SpRY-HF1. Finally, when using SpRY and gRNA A8, no off-targets were cleaved more efficiently than the on-target site in CHANGE-seq experiments (**Sup. Figs. 12** and **13**).

When validating CHANGE-seq nominated off-targets via targeted sequencing in ABE8e-SpRY and gRNA A8 treated fibroblasts, we observed very low levels of off-target editing at 2 of 168 off-target adenines (**Sup. Fig. 14a** and **Supplementary Table 2**), which reached statistical significance only without correcting for multiple hypothesis testing. These off-target adenines are located on chromosome 4 (48561250-48561273 using GRCh38) in an intron of *ZAR1* (mean 0.0171% A-to-G editing in the ABE-treated sample vs. 0.0068% in the naïve control) and on chromosome 7 (16099553-16099576 using GRCh38) in an intron of *CRPPA* (mean 0.0137% A-to-G editing in the ABE-treated sample vs. 0.0058% in the naïve control).

**Supplementary Note 5**: **Comparison of AAV9 and AAV-F capsids.**

To select an AAV serotype for *in vivo* studies, we compared the transduction efficiencies of AAV9 and a recently developed AAV9 capsid variant (AAV-F) that has been shown to mediate efficient delivery to the central nervous system (CNS). Compared to AAV9, AAV-F was described to mediate >170-fold improved transduction of neurons in adult mice after systemic delivery[17,18]. However, the generalizability of AAV-F was not previously assessed via our envisioned *in vivo* approach involving intracerebroventricular (ICV) injections in neonate FVB or SMNΔ7 mice. Thus, we performed experiments involving P1 ICV injections of FVB neonate mice using 2 x $10^{10}$ vg of AAV9-EGFP or AAV-F-EGFP (**Sup. Fig. 16a**). We analyzed EGFP expression in multiple tissues including the brain and spinal cord at 13 days post injection, and observed efficient transduction of both tissues using either serotype assessed via imaging (**Sup. Figs. 16b** and **16c**) or western blots (**Sup. Figs. 16d** and **16e**). While AAV-F-EGFP was more specific for CNS transduction than AAV9-EGFP by exhibiting lower efficiency transduction of other peripheral tissues including liver and heart (**Sup. Fig. 16b-16e**), the overall transduction efficiency of AAV9-EGFP or AAV-F-EGFP in the brain and spinal cord were comparable. Based on these data, we conducted the *in vivo* experiments using AAV9 due to the longstanding use of this serotype academically and clinically, and that our results using AAV-F via ICV injection in neonates were not as striking as those previously observed via systemic injections in adult mice[17]. The lower off-target tissue transduction of AAV-F compared to AAV9 after ICV injection may be useful in studies where expression is restricted to certain tissues. Future studies investigating different delivery methods and other newly engineered AAV capsids with improved CNS transduction[19-21] may improve the delivery of base editors in the CNS, with particular interest in those that can effectively target motor neurons.

**Supplementary Note 6: Details and results of *in vivo* SMNΔ7 mice injections**.

We selected SMNΔ7 mice[22-24] for our study because they have a well-established SMA phenotype and harbor a human copy of the *SMN2* gene, permitting us to assess *in vivo* editing using our human-designed gRNAs from the remainder of our study. The genetic basis of the SMNΔ7 mice[22-24] consists of 3 main mutations in the FVB background. First, exon 2 of the endogenous mouse *Smn* gene was disrupted by the insertion of a lacZ-neo cassette resulting in an in-frame fusion of lacZ. SMNΔ7 mice can be homozygous (Smn+/+ or Smn-/-) or heterozygous (Smn+/-) for the mouse *Smn2 gene.* The Smn+/+ or Smn +/- will not display a phenotype. Homozygous SMNΔ7 mice knocked out for *Smn2* (Smn-/-) develop a severe form of SMA. Second, SMNΔ7 mice carry a copy of the human *SMN2* gene and promoter (fragment of 35.5 kb) integrated into intron 4 of the mouse *Grm7* gene. This inserted *SMN2* is the target of the gRNAs in our present study. Third, SMNΔ7 mice express additional copies of a human *SMN2* cDNA (SMNdelta7) that lacks exon 7, a molecular strategy applied to increase the viability of this model[22]. These additional cDNA copies are not targeted by the gRNAs and base editors in our current study, due to the lack of exon 7.

At 13 days post injections (P13) with N- and C-terminal AAVs to express base editors (**Sup. Fig. 17a**), we observed precise *SMN2* C6T editing in the brain, spinal cord, liver, heart, and skeletal muscle in both SMA (**Sup. Fig. 17b**) or unaffected mice carrying the human *SMN2* (**Sup. Fig. 17c**). The two SMA mice investigated in this experiment weighted 2.7 g and 4.0 g at P13, while the weights of the unaffected mice ranges from 9.2 to 9.9 g at the same time point. Moreover, SMA mice were unable to perform the righting reflex (the motor ability to flip from a supine position), which was performed in under 2 seconds by unaffected mice. Since the lifespans of SMA mice are typically less than 15 days[25], tissues from all mice were collected at P13 for consistency regardless of physical condition of the animals. Due to the rapid and severe phenotype of the SMNΔ7 mouse model, evaluation of potential short-term physiological benefits resulting from base editing is a challenge due to a very narrow therapeutic window (where editing even immediately at P1 may not be sufficient). Thus, although our *SMN2* C6T editing strategy is a potential genetic permanent treatment for SMA, future studies to expand the therapeutic window by delaying disease onset or enabling editing at earlier timepoints (i.e. in utero), may be necessary to extend SMA mouse survival to allow the investigation of longer-term base editing effects.

### Supplementary Figures and Legends

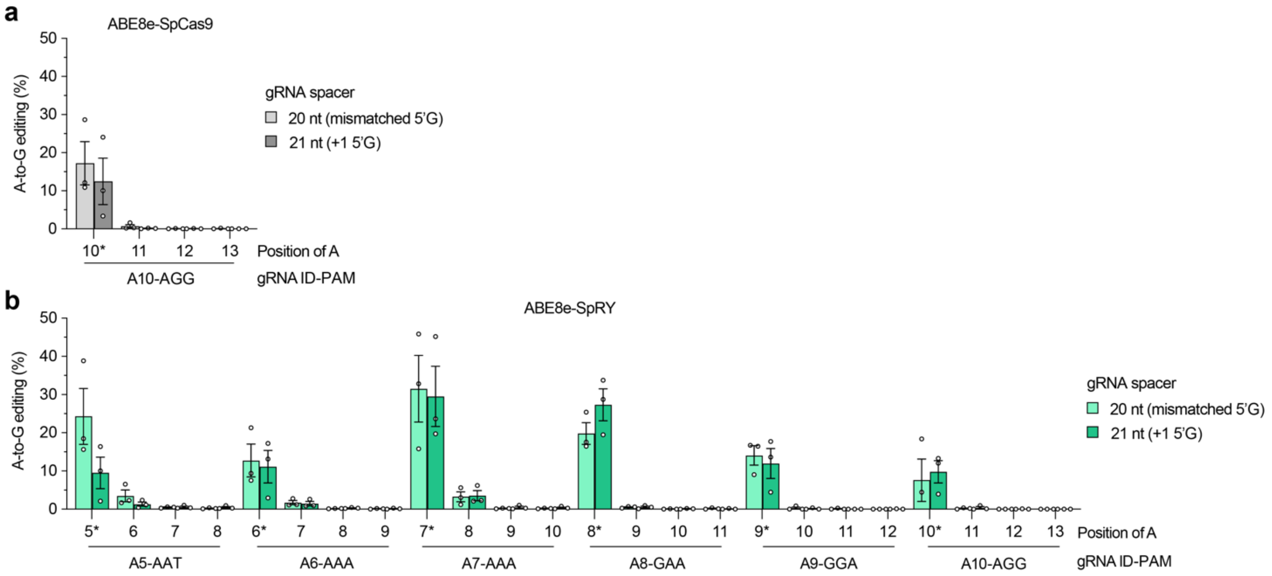


**Supplementary Figure 1. Comparison of 5' gRNA spacer architectures. a,** A-to-G editing of *SMN2* C6T target adenine and other bystander bases (see **Fig. 1b**) when transfecting plasmids encoding gRNAs with 20 nt spacers (harboring matched 5'G or mismatched 5'G bases; see **Supplementary Table 3**) or 21 nt spacers (with an additional +1 5'G base) and ABE8e-SpCas9 (**panel a**) or ABE8e-SpRY (**panel b**) into HEK 293T cells. A-to-G editing assessed by targeted sequencing; mean, s.e.m., and individual datapoints shown for n = 3 independent biological replicates.

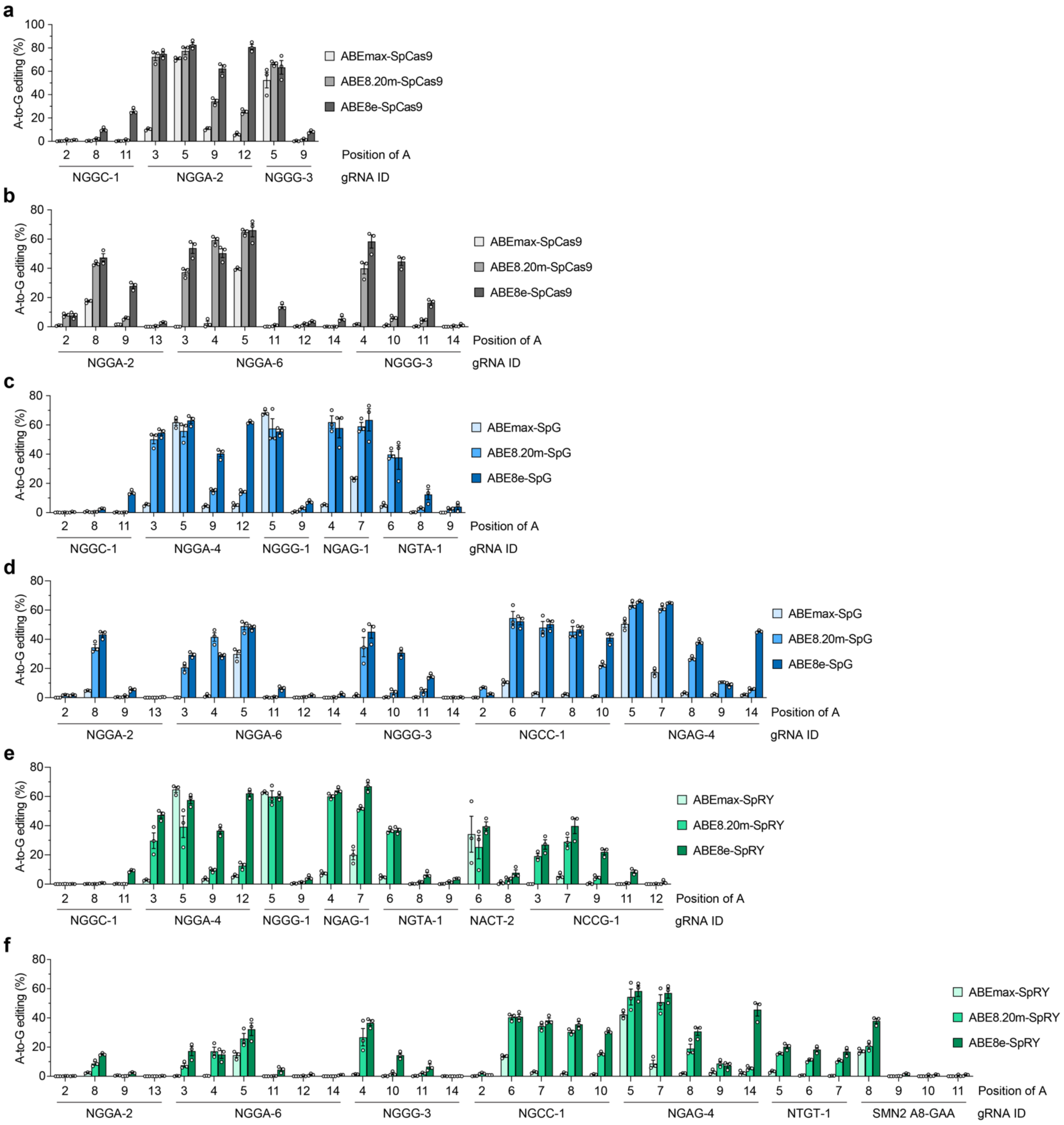


**Supplementary Figure 2. Assessment of ABEs targeting conventional and poly-adenine target sites. a-f,** A-to-G editing of target sites in HEK 293T cells with ABEs harboring deaminase domains ABEmax[10,11], ABE8.20m[12], and ABE8e fused to wild-type SpCas9 (targeting conventional or poly A target sites; **panels a** and **b**, respectively), to SpG[13] (targeting conventional or poly A target sites; **panels c** and **d**, respectively), or to SpRY[13] (targeting conventional or poly A target sites; **panels e** and **f**, respectively). Target sites were selected from previous studies[13,26-28]. A-to-G editing assessed by targeted sequencing; mean, s.e.m., and individual datapoints shown for n = 3 independent biological replicates.

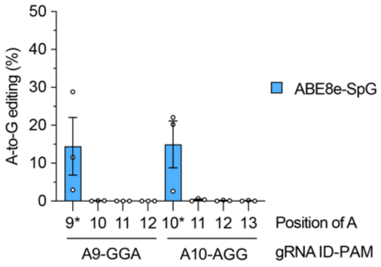


**Supplementary Figure 3. *SMN2* exon 7 editing with ABE8e-SpG.** A-to-G editing in exon 7 of *SMN2* (see **Fig. 1b**) when transfecting plasmids encoding ABE8e-SpG and gRNAs A9 or A10 in HEK 293T cells. A-to-G editing assessed by targeted sequencing; mean, s.e.m., and individual datapoints shown for n = 3 independent biological replicates.

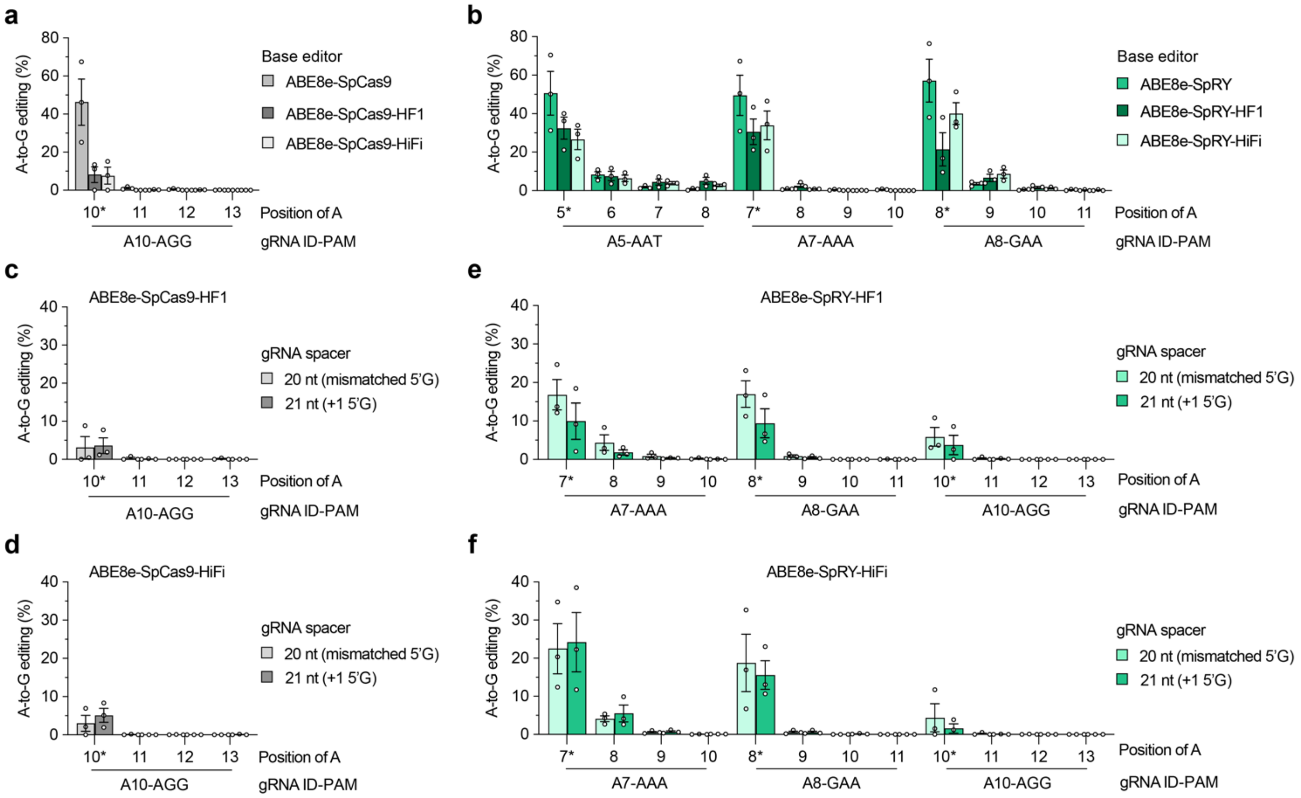


**Supplementary Figure 4. On-target base editing with high-fidelity versions of ABEs**. **a**,**b**, A-to-G editing in exon 7 of *SMN2* (see **Fig. 1b**) when transfecting plasmids encoding gRNA A10 and ABE8e-WT (without or with HF1 or HiFi mutations[4,5]; **panel a**) or gRNAs A5, A7, or A8 with ABE8e-SpRY (without or with HF1 or HiFi mutations; **panel b**) in HEK 293T cells. **c**-**f**, A-to-G editing in exon 7 of *SMN2* when transfecting plasmids encoding gRNAs with 20 nt spacers (harboring matched 5'G or mismatched 5'G bases; see **Supplementary Table 3**) or 21 nt spacers (with an additional +1 5'G base) and ABE8e-SpCas9-HF1 (**panel c**), ABE8e-SpCas9-HiFi (**panel d**), ABE8e-SpRY-HF1 (**panel e**), or ABE8e-WT-HiFi (**panel f**). For **panels a-f**, A-to-G editing assessed by targeted sequencing; mean, s.e.m., and individual datapoints shown for n = 3 independent biological replicates.

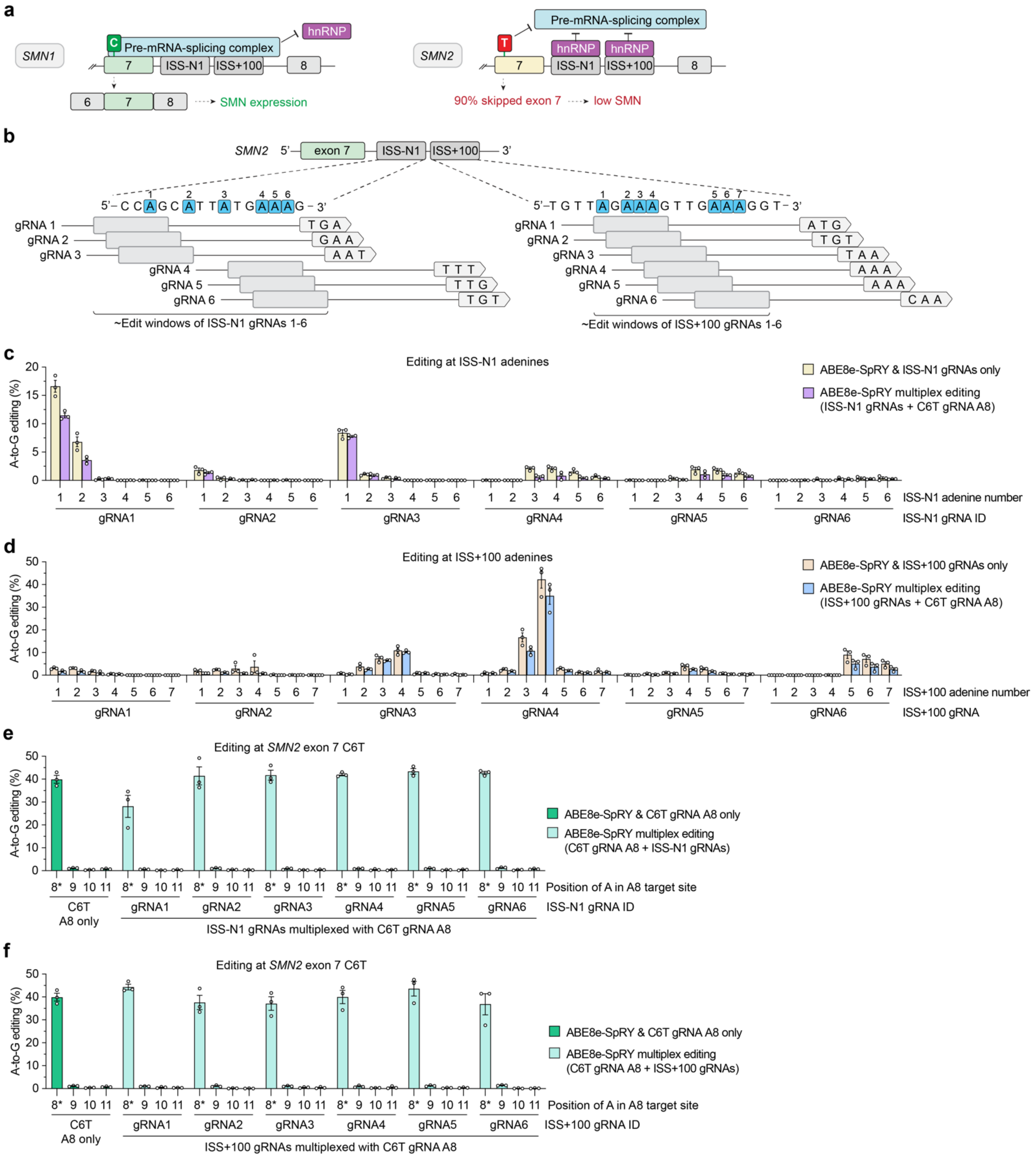


**Supplementary Figure 5. Multiplex base editing of *SMN2* intronic splicing silencers (ISSs). a,** Schematic of ISS binding sites (ISS-N1 and ISS+100) for hnRNP ribonucleoproteins located in *SMN2* intron 7. **b,** Schematic of the ISS-N1 and ISS+100 binding sites, with base editor gRNA target sites and their estimated edit windows. **c,** A-to-G editing of bases in ISS-N1 when transfecting plasmids encoding ABE8e-SpRY and ISS-N1 gRNAs with or without *SMN2* exon 7 gRNA A in HEK 293T cells. **d,** A-to-G editing of bases in ISS+100 when transfecting plasmids encoding ABE8e-SpRY and ISS-N1 gRNAs with or without *SMN2* exon 7 gRNA A in HEK 293T cells.

**e,f,** A-to-G editing of *SMN2* C6T target adenine and other bystander bases when transfecting plasmids encoding ABE8e-SpRY and *SMN2* exon 7 gRNA A8 with or without ISS-N1 gRNAs (**panel e**) or with or without ISS+100 gRNAs (**panel f**) in HEK 293T cells. A-to-G editing assessed by targeted sequencing; mean, s.e.m., and individual datapoints shown for n = 3 independent biological replicates.

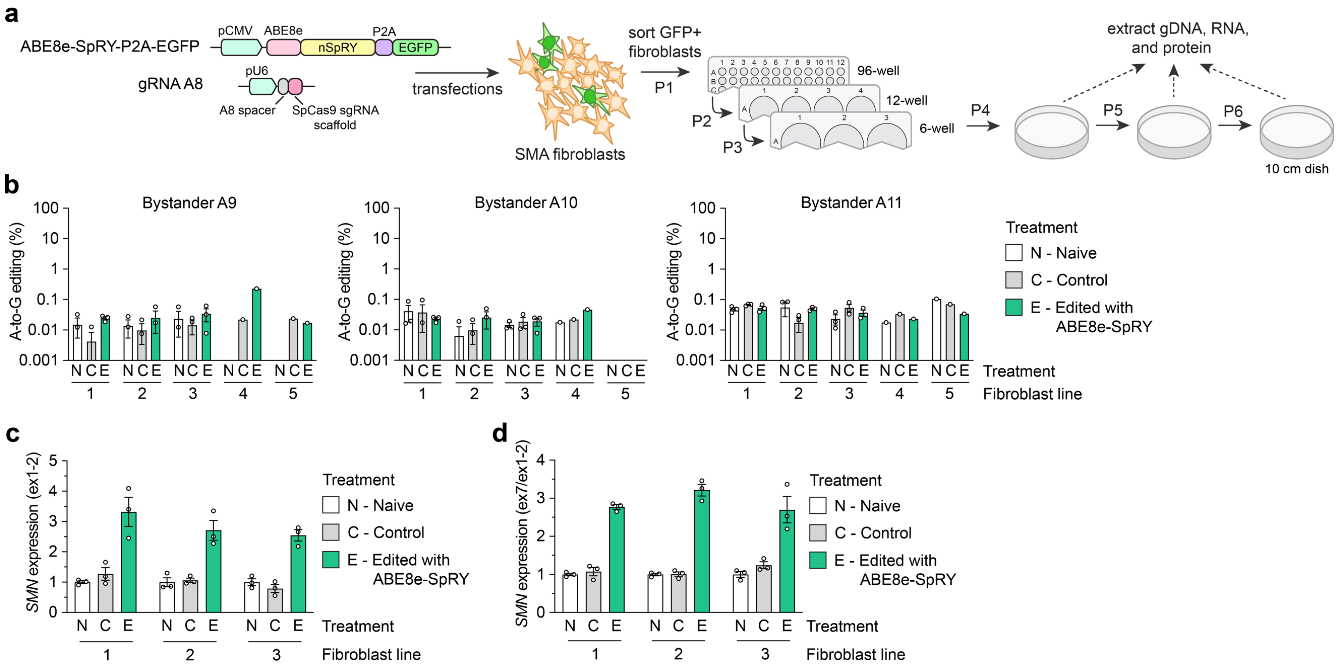


**Supplementary Figure 6. ABE8e-SpRY increases the SMN mRNA expression in SMA patient-derived fibroblasts. a,** Schematic of transfections as well as cell sorting and expansion when performing transfections using plasmids encoding ABE8e-SpRY and gRNA A8 in SMA patient-derived fibroblasts. **b**, A-to-G editing of bystander adenines adjacent to *SMN2* C6T when transfecting plasmids encoding ABE8e-SpRY and *SMN2* exon 7 gRNA A8. A-to-G editing assessed by targeted sequencing; mean, s.e.m., and individual datapoints shown. **c**, SMN mRNA expression measured across the exon 1 and 2 junction (for both *SMN1* and *SMN2*) normalized by GAPDH, assessed by ddPCR for each of the three edited SMA fibroblast lines. **d,** *SMN2* exon 7 mRNA expression normalized by the *SMN1* and *SMN2* exon 1-2 junction. mRNA expression assessed across three edited SMA fibroblast lines. For all data in **panels b-d**, GFP-positive fibroblasts were sorted after transfections and grew in cultures for 6 passages prior to freezing (see **panel a**); genomic DNA, RNA, and protein samples were collected from three independent passages (P4-P6) for assays.

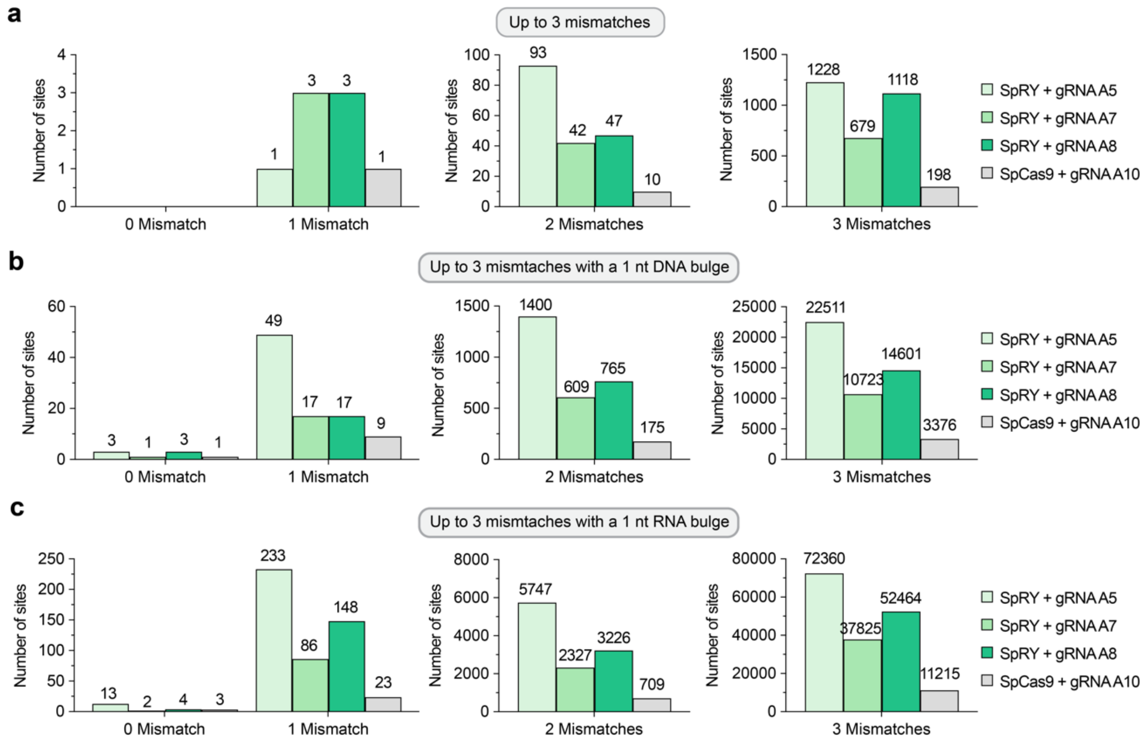


**Supplementary Figure 7. *In silico* annotated off-target sites. a-c**, Putative off-target sites were identified computationally using CasOFFinder[29] for WT SpCas9 with gRNA A10, and for SpRY with gRNAs A5, A7, and A8. Off-targets were enumerated when considering up to 3 mismatches (**panel a**), 3 mismatches with a 1 nt DNA bulge (**panel b**), or 3 mismatches with a 1 nt RNA bulge (**panel c**).

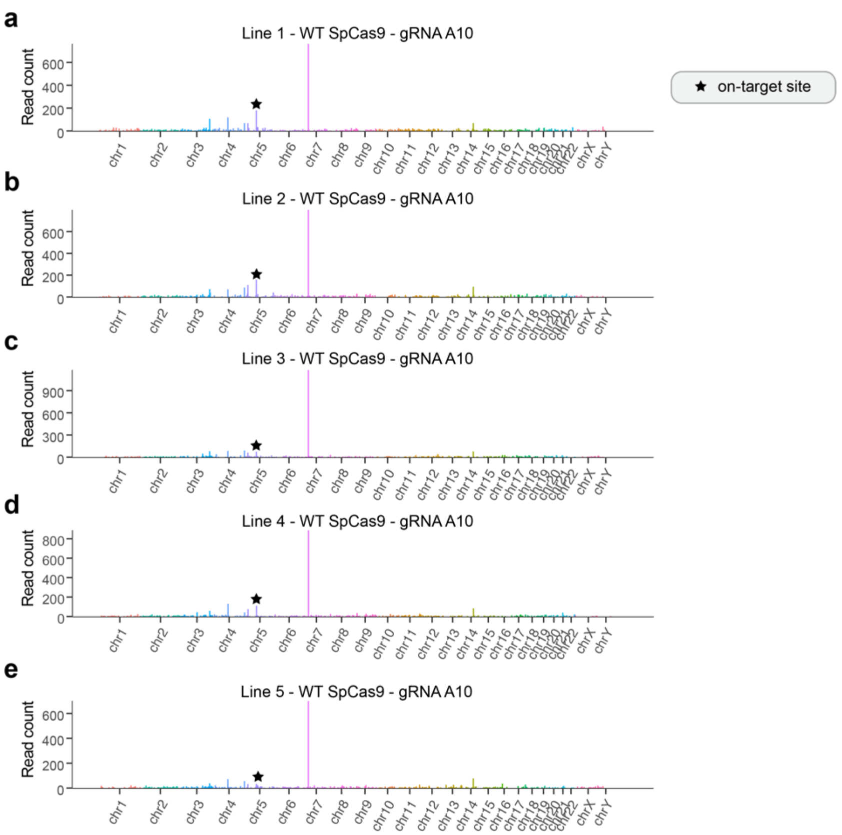


**Supplementary Figure 8. Locations of CHANGE-seq detected sites for WT SpCas9 and gRNA A10. a-e,** Manhattan plots of CHANGE-seq-detected on- and off-target sites, ordered by chromosomal position with bar heights proportional to CHANGE-seq read counts. Experiments were performed with WT SpCas9 nuclease and gRNA A10 along with genomic DNA from SMA patient-derived fibroblast lines 1-5 (**panels a-e**, respectively). The on-target site is indicated using a black star.

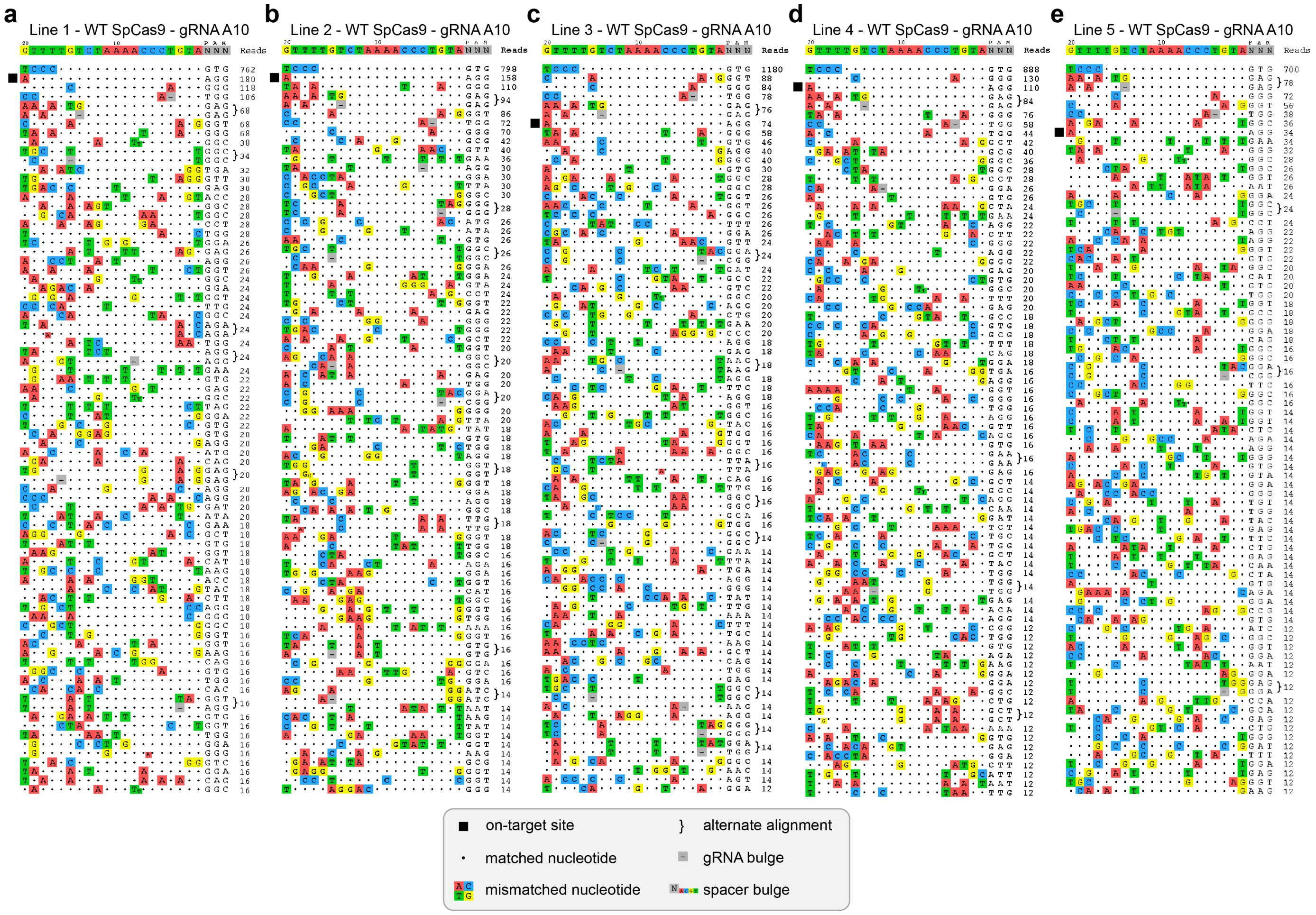


**Supplementary Figure 9. CHANGE-seq detected off-target sites for WT SpCas9 and gRNA A10**. **a-e**, Rank-ordered visualization of the top approximately 80 on- and off-target genomic sites identified by CHANGE-seq for WT-SpCas9 nuclease and gRNA A10, using genomic DNA from SMA patient-derived fibroblast lines 1-5 (**panels a-e**, respectively). The on-target site is indicated using a black box; alternate alignments are shown for sites that are potentially targeted via 1 nt DNA or gRNA spacer bulges.

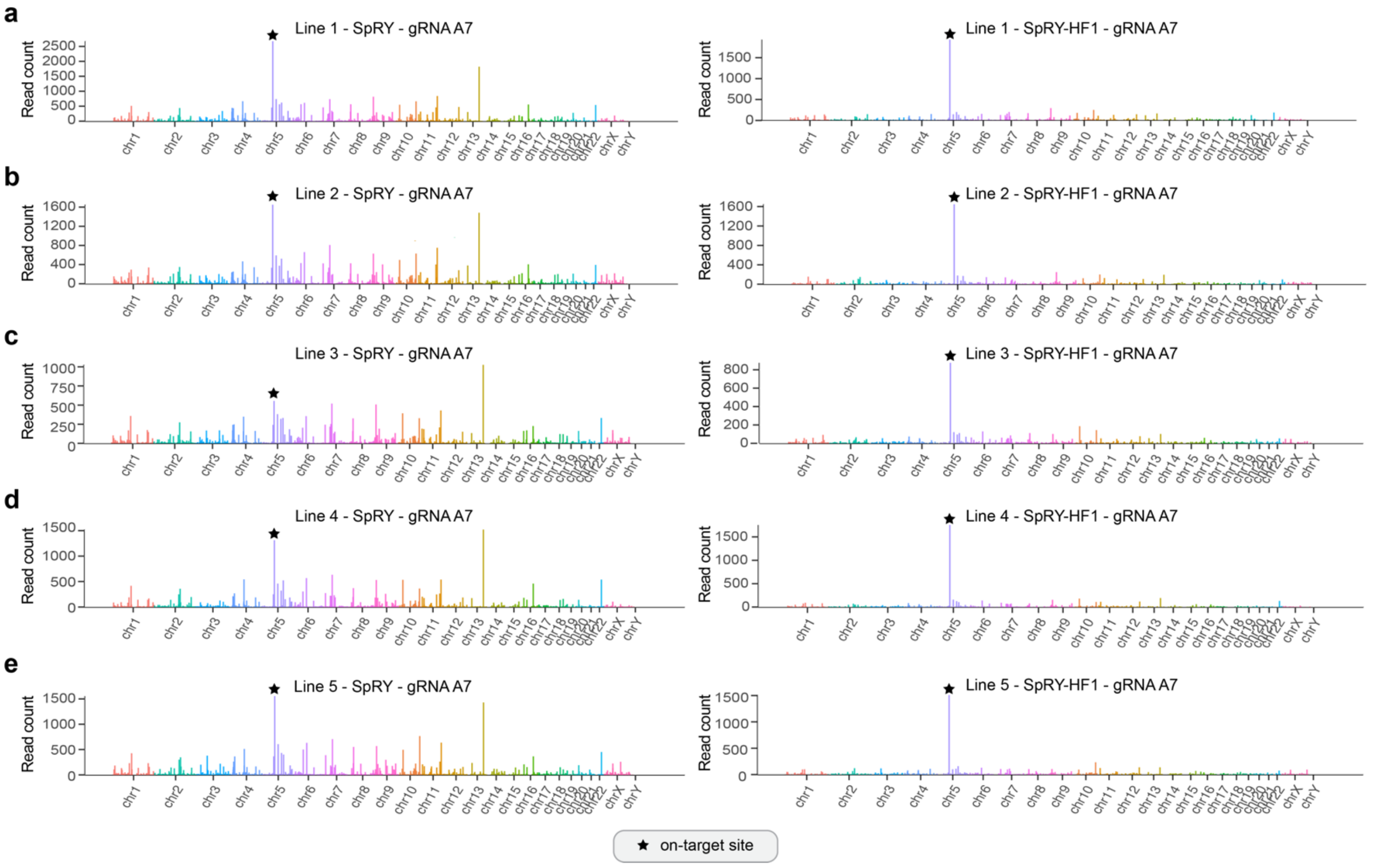


**Supplementary Figure 10. Locations of CHANGE-seq detected sites for SpRY and gRNA A7. a-e,** Manhattan plots of CHANGE-seq-detected on- and off-target sites, ordered by chromosomal position with bar heights proportional to CHANGE-seq read counts. Experiments were performed with SpRY nuclease[13] (left panels) or SpRY-HF1[5,13] nuclease (right panels) and gRNA A7 along with genomic DNA from SMA patient-derived fibroblast lines 1-5 (**panels a-e**, respectively). The on-target site is indicated using a black star.

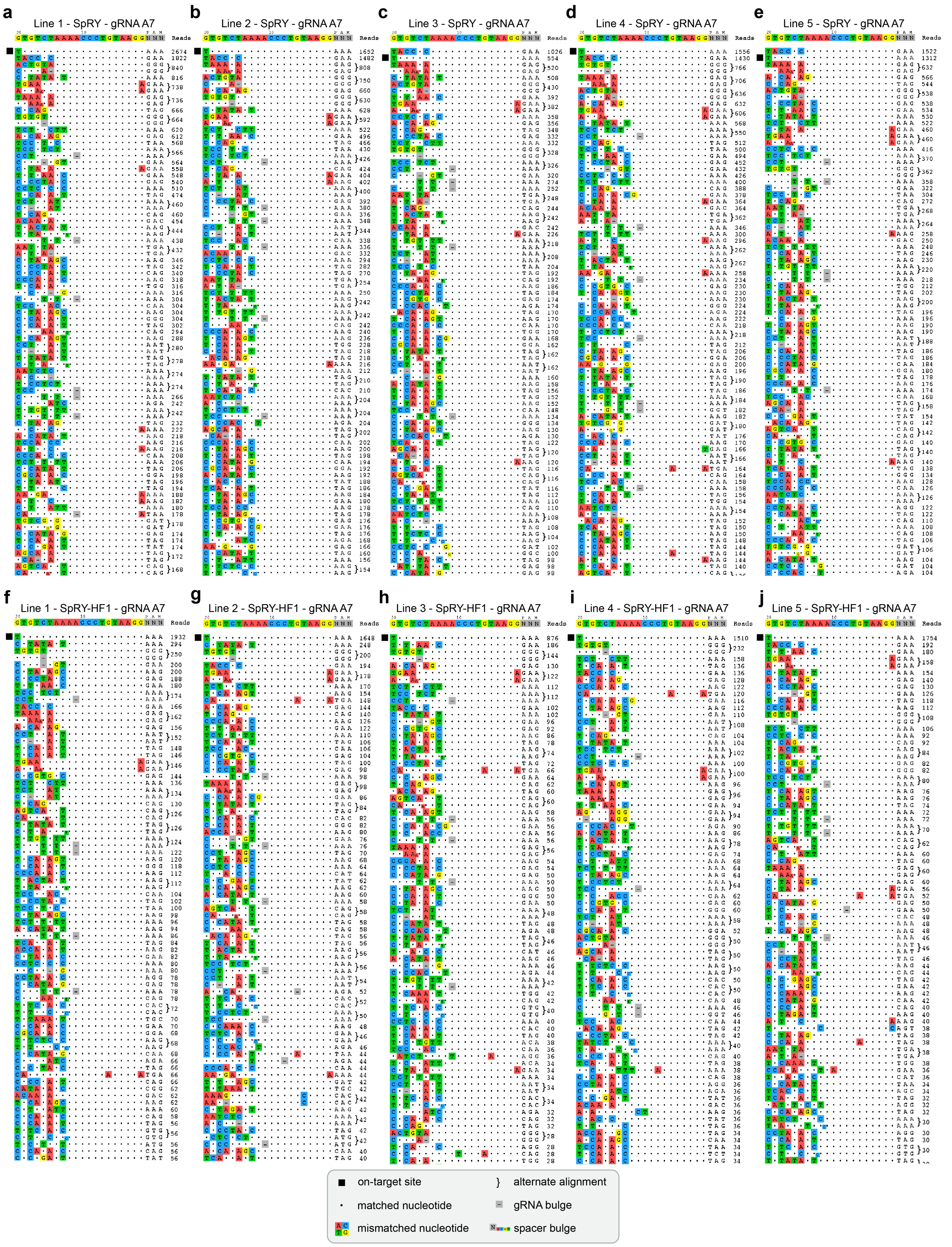
a
Line 1 - SpRY - gRNA A7
b
Line 2 - SpRY - gRNA A7
c
Line 3 - SpRY - gRNA A7
d
Line 4 - SpRY - gRNA A7
e
Line 5 - SpRY - gRNA A7
f
Line 1 - SpRY-HF1 - gRNA A7
g
Line 2 - SpRY-HF1 - gRNA A7
h
Line 3 - SpRY-HF1 - gRNA A7
i
Line 4 - SpRY-HF1 - gRNA A7
j
Line 5 - SpRY-HF1 - gRNA A7
GTGTCTAAAACCCTGTAAGGNNN
PAM
Reads
on-target site
alternate alignment
matched nucleotide
gRNA bulge
mismatched nucleotide
spacer bulge

**Supplementary Figure 11. CHANGE-seq detected off-target sites for SpRY or SpRY-HF1 with gRNA A7**. **a-j**, Rank-ordered visualization of the top approximately 80 on- and off-target genomic sites identified by CHANGE-seq for SpRY nuclease (**panels a-e**) or SpRY-HF1 nuclease (**panels f-j**) and gRNA A7, using genomic DNA from SMA patient-derived fibroblast lines 1-5, respectively. The on-target site is indicated using a black box; alternate alignments are shown for sites that are potentially targeted via 1 nt DNA or gRNA spacer bulges.

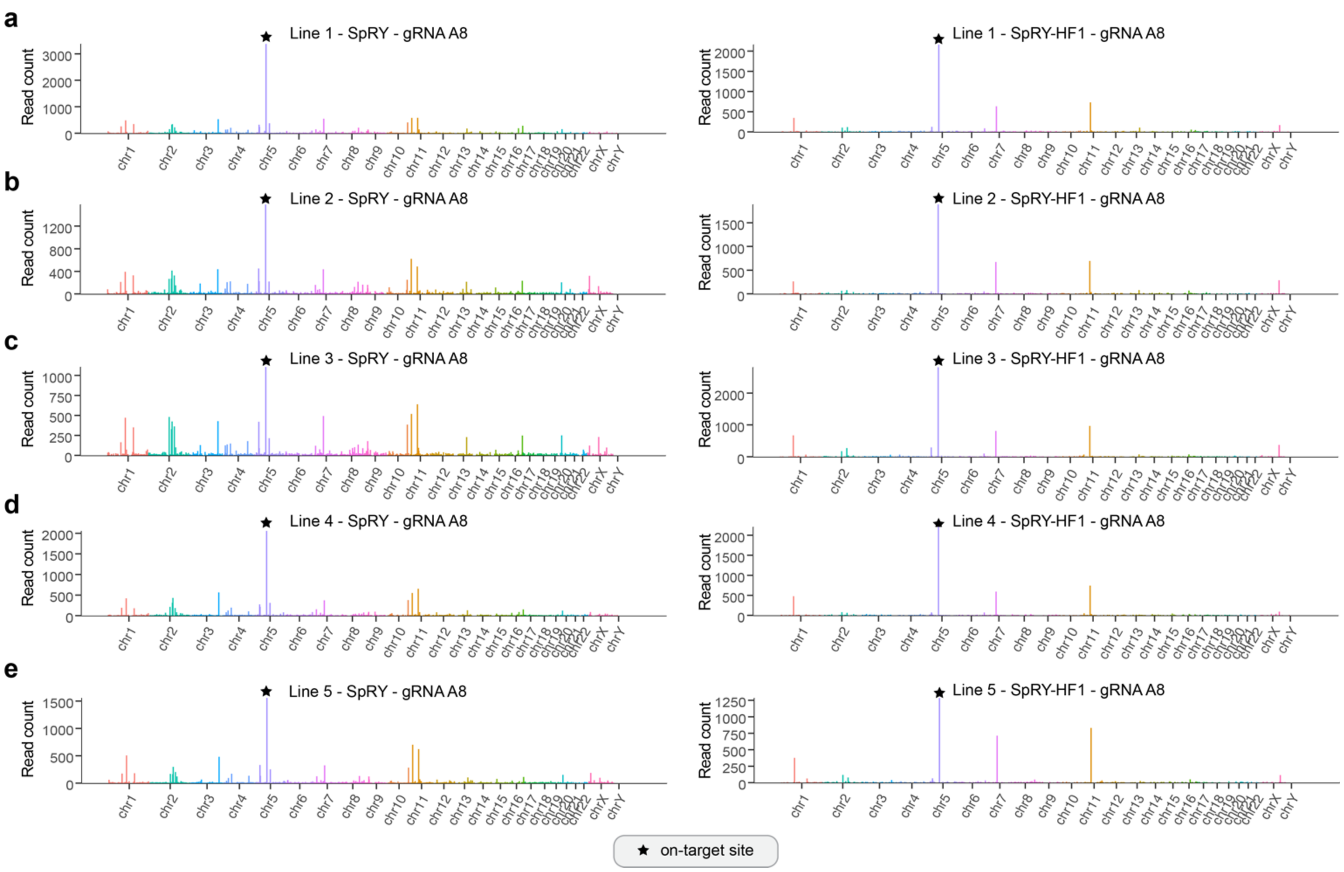


**Supplementary Figure 12. Locations of CHANGE-seq detected sites for SpRY and gRNA A8. a-e,** Manhattan plots of CHANGE-seq-detected on- and off-target sites, ordered by chromosomal position with bar heights proportional to CHANGE-seq read counts. Experiments were performed with SpRY nuclease (left panels) or SpRY-HF1 nuclease (right panels) and gRNA A8 along with genomic DNA from SMA patient-derived fibroblast lines 1-5 (**panels a-e**, respectively). The on-target site is indicated using a black star.

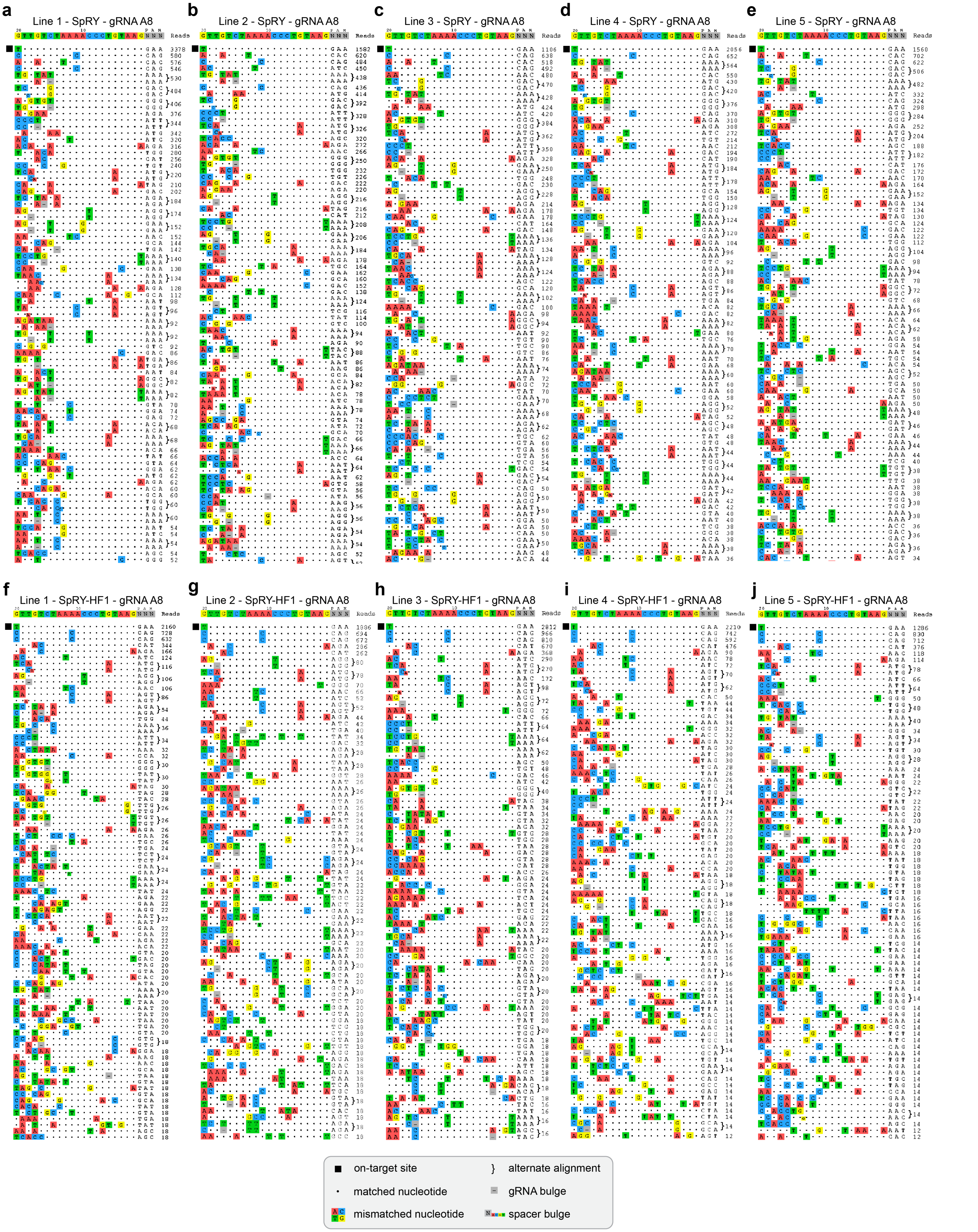
a Line 1 - SpRY - gRNA A8
b Line 2 - SpRY - gRNA A8
c Line 3 - SpRY - gRNA A8
d Line 4 - SpRY - gRNA A8
e Line 5 - SpRY - gRNA A8
f Line 1 - SpRY-HF1 - gRNA A8
g Line 2 - SpRY-HF1 - gRNA A8
h Line 3 - SpRY-HF1 - gRNA A8
i Line 4 - SpRY-HF1 - gRNA A8
j Line 5 - SpRY-HF1 - gRNA A8
Reads
PAM
on-target site
} alternate alignment
matched nucleotide
gRNA bulge
mismatched nucleotide
spacer bulge

**Supplementary Figure 13. CHANGE-seq detected off-target sites for SpRY or SpRY-HF1 with gRNA A7**. **a-j**, Rank-ordered visualization of the top approximately 80 on- and off-target genomic sites identified by CHANGE-seq for SpRY nuclease (**panels a-e**) or SpRY-HF1 nuclease (**panels f-j**) and gRNA A7, using genomic DNA from SMA patient-derived fibroblast lines 1-5, respectively. The on-target site is indicated using a black box; alternate alignments are shown for sites that are potentially targeted via 1 nt DNA or gRNA spacer bulges.

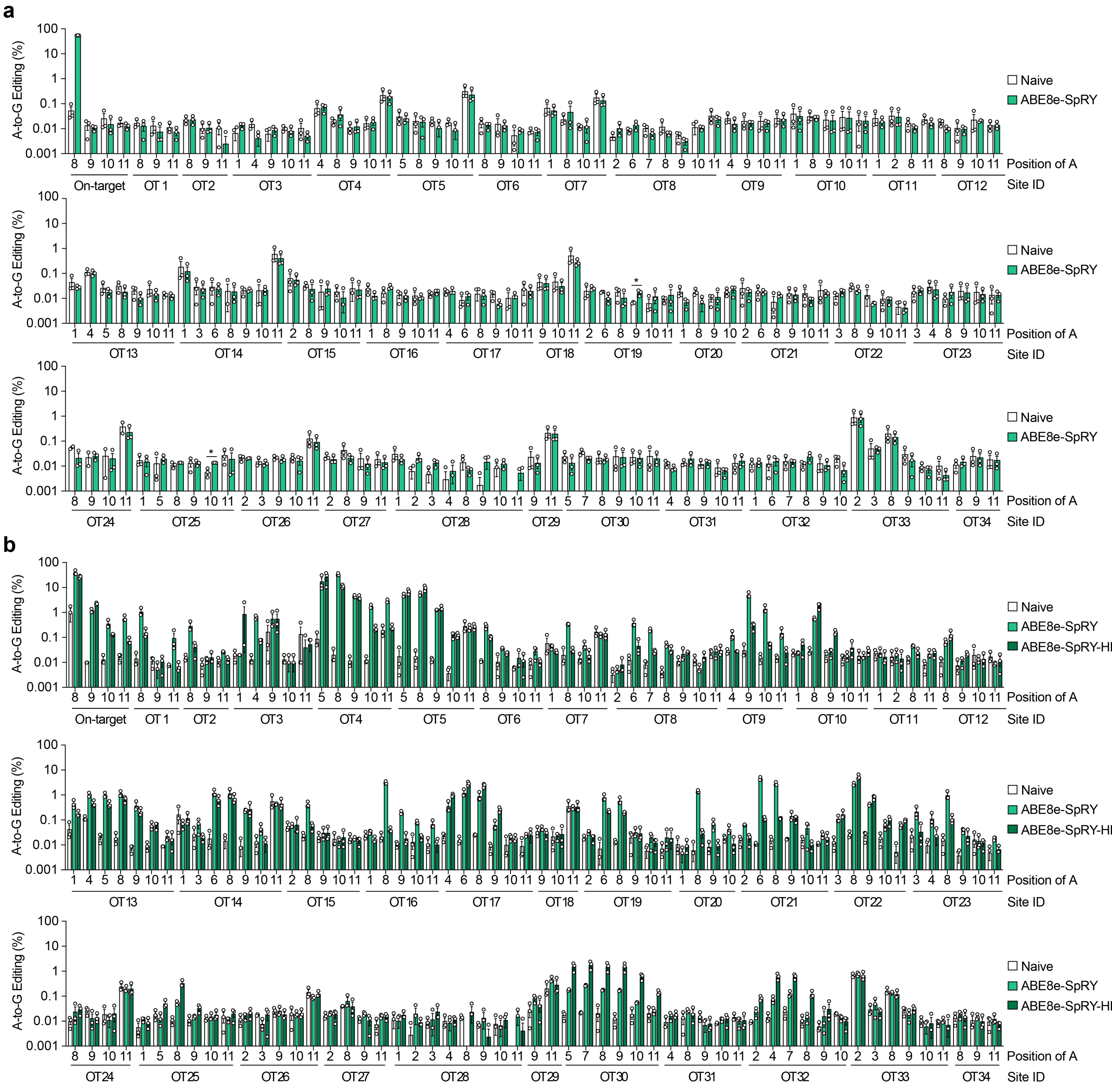


**Supplementary Figure 14. Analysis of editing at CHANGE-seq nominated sites. a,b,** Assessement of A-to-G base editing at the on-target and top 34 off-target sites selected from CHANGE-seq datasets, performed via targeted sequencing of genomic DNA from SMA fibroblast lines 1, 2, and 3 treated with ABE8e-SpRY and gRNA A8 (each datapoint for n = 3 is from a different fibroblast line; **panel a**), or sequencing of genomic DNA from HEK293T cells treated with ABE8e-SpRY or ABE8e-SpRY-HF1 and gRNA A8 (each datapoint for n = 3 is from an independent biological replicate transfection; **panel b**). Mean, s.e.m, and individual datapoints shown for n = 3 as described above; the asterisks in **panel a** indicate adenines with a statistically significant difference between Naïve and ABE8e-SpRY edited samples. **P* = 0.0155 (OT19) or 0.0039 (OT25), two-tailed Student's t-test. Adenines with a statistically significant differences in **panel b** are presented in **Supplementary Table 2**.

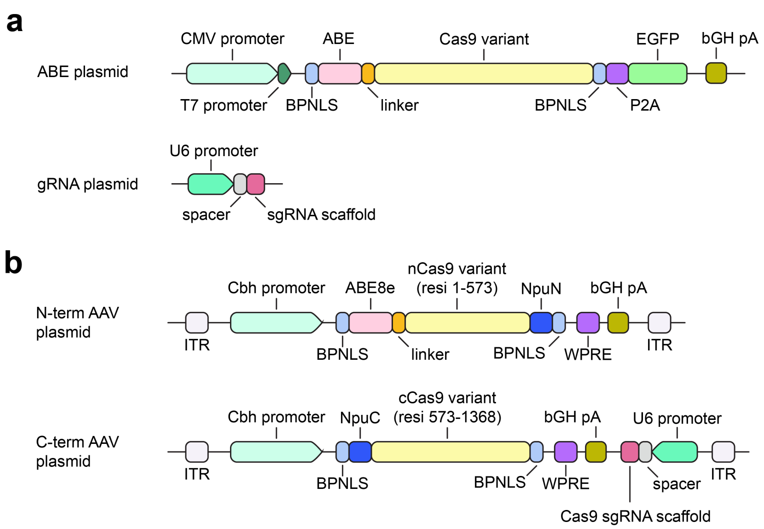


**Supplementary Figure 15. ABE and gRNA plasmids used for transfections or AAV production.** **a**, Schematic of the plasmids used for conventional transfections, with one expressing the ABE enzyme and the other expressing the gRNA. **b**, Schematic of the plasmids used for AAV production, which encode the ABE8e deaminase domain and an N-terminal fragment of Cas9 in the N-term AAV plasmids, and the C-terminal fragment of Cas9 and the gRNA expression cassette in the C-term plasmid. The N- and C-terminal fragments of Cas9 are joined post-translationally via the Npu intein[30].

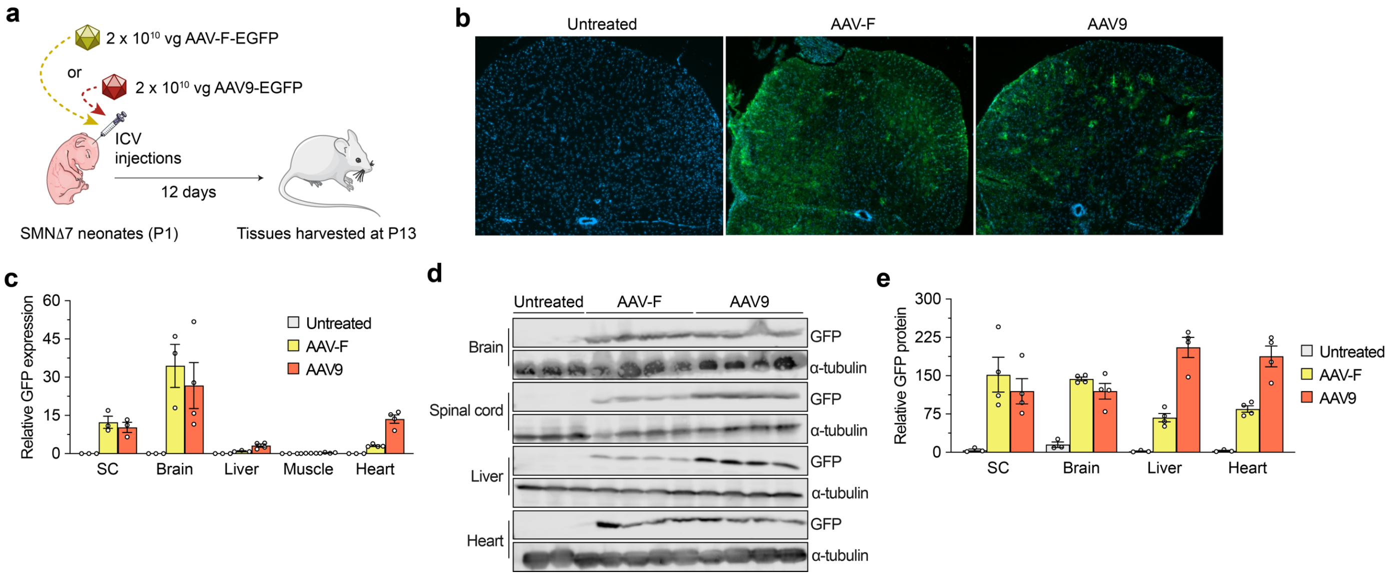


**Supplementary Figure 16. Comparison of the transduction efficiencies of AAV9 and AAV-F in mice.** **a**, Schematic of P1 intracerebroventricular (ICV) injections in FVB mice with 2 x $10^{10}$ vg of AAV9 or AAV-F vectors that express EGFP. **b**, Representative immunofluorescence images to illustrate GFP expression (green) and nuclei staining (dapi; blue) in P13 sections of the spinal cords of P1 FVB mice injected ICV with AAV9-EGFP or AAV-F-EGFP. **c,** Quantification of EGFP mRNA expression, assessed by ddPCR and normalized by GAPDH expression in different tissues at 13 days post injection. **d-e,** Representative images and quantification (**panels d** and **e**, respectively) of immunoblots probing for GFP (~27 kDa) and alpha-tubulin (~50 kDa) in different tissues at 13 days post injection. n = 3 untreated mice and 4 treated mice for either AAV9 or AAV-F group.

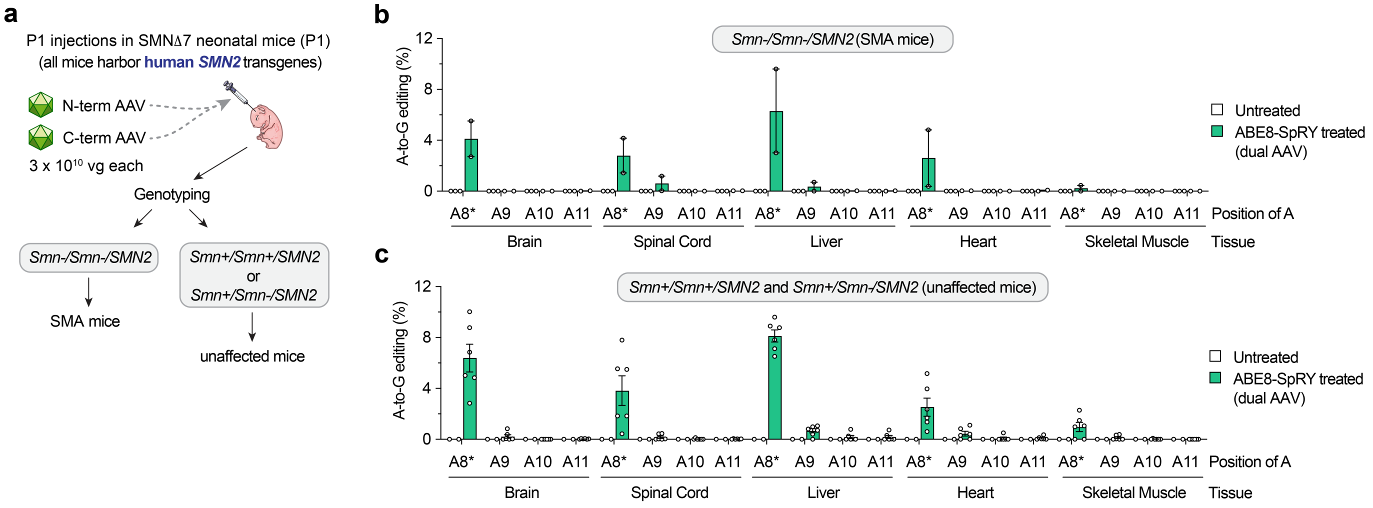


**Supplementary Figure 17. *SMN2* C6T base editing plotted by SMNΔ7 mouse genotype. a**, Schematic of injections and genotyping to segregate mice into SMA (homozygous null for Smn) or unaffected mice (heterozygous or wild-type). Note that all mice are transgenic for human *SMN2*. **b,c,** A-to-G editing of *SMN2* C6T target adenine and other bystander bases divided into SMA mice ($Smn^{-/-}$ with n = 2 mice) (**panel b**), or unaffected mice ($Smn^{+/-}$ and $Smn^{+/+}$ with n = 4 mice and 1 mouse, respectively) (**panel c**).

### Supplementary References


1 Ma, H. *et al.* Pol III Promoters to Express Small RNAs: Delineation of Transcription Initiation. *Mol Ther Nucleic Acids* **3**, e161, doi:10.1038/mtna.2014.12 (2014).

2 Gao, Z., Harwig, A., Berkhout, B. & Herrera-Carrillo, E. Mutation of nucleotides around the +1 position of type 3 polymerase III promoters: The effect on transcriptional activity and start site usage. *Transcription* **8**, 275-287, doi:10.1080/21541264.2017.1322170 (2017).

3 Kim, S., Bae, T., Hwang, J. & Kim, J. S. Rescue of high-specificity Cas9 variants using sgRNAs with matched 5' nucleotides. *Genome Biol* **18**, 218, doi:10.1186/s13059-017-1355-3 (2017).

4 Vakulskas, C. A. *et al.* A high-fidelity Cas9 mutant delivered as a ribonucleoprotein complex enables efficient gene editing in human hematopoietic stem and progenitor cells. *Nat Med* **24**, 1216-1224, doi:10.1038/s41591-018-0137-0 (2018).

5 Kleinstiver, B. P. *et al.* High-fidelity CRISPR-Cas9 nucleases with no detectable genome-wide off-target effects. *Nature* **529**, 490-495, doi:10.1038/nature16526 (2016).

6 Zhang, D. *et al.* Perfectly matched 20-nucleotide guide RNA sequences enable robust genome editing using high-fidelity SpCas9 nucleases. *Genome Biol* **18**, 191, doi:10.1186/s13059-017-1325-9 (2017).

7 He, X. *et al.* Boosting activity of high-fidelity CRISPR/Cas9 variants using a tRNA(Gln)-processing system in human cells. *J Biol Chem* **294**, 9308-9315, doi:10.1074/jbc.RA119.007791 (2019).

8 Richter, M. F. *et al.* Phage-assisted evolution of an adenine base editor with improved Cas domain compatibility and activity. *Nat Biotechnol* **38**, 883-891, doi:10.1038/s41587-020-0453-z (2020).

9 Arbab, M. *et al.* Determinants of Base Editing Outcomes from Target Library Analysis and Machine Learning. *Cell* **182**, 463-480 e430, doi:10.1016/j.cell.2020.05.037 (2020).

10 Gaudelli, N. M. *et al.* Programmable base editing of A*T to G*C in genomic DNA without DNA cleavage. *Nature* **551**, 464-471, doi:10.1038/nature24644 (2017).

11 Koblan, L. W. *et al.* Improving cytidine and adenine base editors by expression optimization and ancestral reconstruction. *Nat Biotechnol* **36**, 843-846, doi:10.1038/nbt.4172 (2018).

12 Gaudelli, N. M. *et al.* Directed evolution of adenine base editors with increased activity and therapeutic application. *Nat Biotechnol* **38**, 892-900, doi:10.1038/s41587-020-0491-6 (2020).

13 Walton, R. T., Christie, K. A., Whittaker, M. N. & Kleinstiver, B. P. Unconstrained genome targeting with near-PAMless engineered CRISPR-Cas9 variants. *Science* **368**, 290-296, doi:10.1126/science.aba8853 (2020).

14 Christie, K. A. & Kleinstiver, B. P. Making the cut with PAMless CRISPR-Cas enzymes. *Trends Genet* **37**, 1053-1055, doi:10.1016/j.tig.2021.09.002 (2021).

15 Newby, G. A. *et al.* Base editing of haematopoietic stem cells rescues sickle cell disease in mice. *Nature* **595**, 295-302, doi:10.1038/s41586-021-03609-w (2021).

16 Lazzarotto, C. R. *et al.* CHANGE-seq reveals genetic and epigenetic effects on CRISPR-Cas9 genome-wide activity. *Nat Biotechnol* **38**, 1317-1327, doi:10.1038/s41587-020-0555-7 (2020).

17 Hanlon, K. S. *et al.* Selection of an Efficient AAV Vector for Robust CNS Transgene Expression. *Mol Ther Methods Clin Dev* **15**, 320-332, doi:10.1016/j.omtm.2019.10.007 (2019).

18 Huang, Q. *et al.* Targeting AAV vectors to the CNS via <em>de novo</em> engineered capsid-receptor interactions. *bioRxiv*, 2022.2010.2031.514553, doi:10.1101/2022.10.31.514553 (2022).

19 Goertsen, D. *et al.* AAV capsid variants with brain-wide transgene expression and decreased liver targeting after intravenous delivery in mouse and marmoset. *Nat Neurosci* **25**, 106-115, doi:10.1038/s41593-021-00969-4 (2022).

20 Ravindra Kumar, S. *et al.* Multiplexed Cre-dependent selection yields systemic AAVs for targeting distinct brain cell types. *Nat Methods* **17**, 541-550, doi:10.1038/s41592-020-0799-7 (2020).

21 Stanton, A. C. *et al.* Systemic administration of novel engineered AAV capsids facilitates enhanced transgene expression in the macaque CNS. *Med (N Y)* **4**, 31-50 e38, doi:10.1016/j.medj.2022.11.002 (2023).

22 Le, T. T. *et al.* SMNDelta7, the major product of the centromeric survival motor neuron (SMN2) gene, extends survival in mice with spinal muscular atrophy and associates with full-length SMN. *Hum Mol Genet* **14**, 845-857, doi:10.1093/hmg/ddi078 (2005).

23 Lutz, C. M. *et al.* Postsymptomatic restoration of SMN rescues the disease phenotype in a mouse model of severe spinal muscular atrophy. *J Clin Invest* **121**, 3029-3041, doi:10.1172/JCI57291 (2011).

24 Kray, K. M., McGovern, V. L., Chugh, D., Arnold, W. D. & Burghes, A. H. M. Dual SMN inducing therapies can rescue survival and motor unit function in symptomatic Δ7SMA mice. *Neurobiol Dis* **159**, 105488, doi:10.1016/j.nbd.2021.105488 (2021).
25 Buettner, J. M. *et al.* Central synaptopathy is the most conserved feature of motor circuit pathology across spinal muscular atrophy mouse models. *iScience* **24**, 103376, doi:10.1016/j.isci.2021.103376 (2021).
26 Nishimasu, H. *et al.* Engineered CRISPR-Cas9 nuclease with expanded targeting space. *Science* **361**, 1259-1262, doi:10.1126/science.aas9129 (2018).
27 Hu, J. H. *et al.* Evolved Cas9 variants with broad PAM compatibility and high DNA specificity. *Nature* **556**, 57-63, doi:10.1038/nature26155 (2018).
28 Kleinstiver, B. P. *et al.* Engineered CRISPR-Cas9 nucleases with altered PAM specificities. *Nature* **523**, 481-485, doi:10.1038/nature14592 (2015).
29 Bae, S., Park, J. & Kim, J. S. Cas-OFFinder: a fast and versatile algorithm that searches for potential off-target sites of Cas9 RNA-guided endonucleases. *Bioinformatics* **30**, 1473-1475, doi:10.1093/bioinformatics/btu048 (2014).
30 Levy, J. M. *et al.* Cytosine and adenine base editing of the brain, liver, retina, heart and skeletal muscle of mice via adeno-associated viruses. *Nat Biomed Eng* **4**, 97-110, doi:10.1038/s41551-019-0501-5 (2020).
